# Lysine lactylation regulates ATF4-mediated stress responses under glucose starvation in canine hemangiosarcoma

**DOI:** 10.1101/2025.08.12.669864

**Authors:** Tamami Suzuki, Kazuki Heishima, Jumpei Yamazaki, Masaya Yamazaki, Ryohei Kinoshita, Sangho Kim, Kenji Hosoya, Yuko Okamatsu-Ogura, Michihito Sasaki, Peng Xu, Qin Yan, Takashi Kimura, Keisuke Aoshima

## Abstract

Hemangiosarcoma (HSA) is a malignant endothelial tumor that occurs frequently in dogs but is rare in other species including humans. Due to its aggressive behavior and limited therapeutic options, patient prognosis is generally poor. Tumor cells produce excess lactate via anerobic glycolysis, and it regulate gene expressions through histone lactylation in response to cellular metabolic conditions. However, how histone lactylation affects biological behavior under glucose-limited conditions in HSA remains unknown. Here, we established canine HSA cell lines and patient-derived xenograft models and investigated the role of histone lactylation during glucose deprivation. HSA cells exhibited higher global histone lactylation levels than normal endothelial cells. Although glucose restriction reduced global histone lactylation levels, Cleavage Under Targets and Tagmentation (CUT&Tag) analysis revealed enrichment of lactylation peaks at transcription-start sites (TSSs) of ATF4-regulated stress-response, asparagine biosynthesis and immune-related genes. TSSs of stress-response genes were co-occupied with RNA polymerase II phosphorylated at serine 5 and showed increased gene expressions, suggesting that lactylation at TSSs activated transcription under glucose-deprived conditions. [U-^13^C]glutamine tracing indicated that HSA cells synthesized asparagine from glutamine when glucose was scarce. Asparagine supplementation modestly activated cell proliferation. In HSA patient tissues, H3K18la levels were heterogeneous, and M2-like macrophages preferentially infiltrated tumor regions showing low histone lactylation levels. Consistently, glucose-starved HSA cells attracted macrophages and induced M2-like polarization *in vitro*. These findings demonstrate that lysine lactylation, possibly histone lactylation, persists even under glucose-deprived conditions and regulate transcription that supports tumor cell survival and fosters a pro-tumor microenvironment.

**One Sentence Summary:** Lysine lactylation is enriched at TSSs of stress-response genes under glucose starvation and associated with their transcription in canine hemangiosarcoma.

## INTRODUCTION

Hemangiosarcoma (HSA) is a malignant tumor of vascular endothelial cells. Although it has been reported in domestic and wildlife animals, dogs have a disproportionately high risk of HSA development (*1–4*) In dogs, HSA accounts for 1.3–2.8% of all canine tumors and often arise in the spleen, liver and the right atrium of the heart (*5–8*). One study reported that 4,997 HSA cases in 2.2 million dogs (2,271 dogs per one million dogs) (*9*). Given that HSA is highly invasive, rapid metastases and tumor rupture are frequently accompanied and contribute to the poor prognosis (*10*). While surgery and doxorubicin-based chemotherapy remain the standard of care, median survival times are only about five to six months and the one-year survival rate for dogs is less than 16% (*11*). This is because limited basic research tools and knowledge of molecular pathogenesis have hindered the development of a novel effective therapeutic. HSA shares noticeable similarities to human angiosarcoma in morphological features, genetic mutations (e.g., *TP53*, *PIK3CA*, *ATRX*), and aggressive clinical behavior (*12–14*). Human angiosarcoma is extremely rare, and several studies have estimated an incidence of ∼ 3 cases and ∼1.5–2.6 cases per one million person in the United States and Europe, respectively (*15, 16*). Due to this rarity, canine HSA is recognized as a valuable spontaneous model for studying angiosarcoma biology and evaluating novel therapeutics (*13, 17*).

Like many cancers, HSA cells are expected to reprogram their metabolism for rapid proliferation. Tumor cells exhibit metabolic profiles distinct from those of normal cells, enabling them to generate sufficient ATP and nucleotide building blocks. These changes allow tumor cells to survive in harsh environments such as hypoxia and poor nutrient conditions (*18, 19*). Even in anerobic conditions, tumor cells drive glycolysis on top of oxidate phosphorylation (OXPHOS), which results in excess lactate production (*20*). Glutamine is used to supply nucleotides and intermediate metabolites via anaplerosis (*21, 22*), while it also contributes to lactate production via glutaminolysis (*23*). Recent works have shown that lactate is not merely a waste product, rather it is an important substrate for an epigenetic mark, histone lysine lactylation (*24*). In macrophages, histone lactylation upregulates reparative genes such as *Arg1*, and its levels are reduced by glycolysis inhibition and rescued by exogenous lactate supplementation (*25*). In tumors, histone lactylation has been implicated in promoting immunosuppression in glioblastoma and ocular melanoma and it is contributed to tumor progression (*26–30*). Despite the increased number of histone-lactylation studies, most have been conducted under high-lactate conditions. Little is known about the roles of histone lactylation under low lactate conditions.

Furthermore, histone lactylation study has not yet been reported in HSA and human angiosarcoma. Actively proliferating normal endothelial cells rely almost exclusively on aerobic glycolysis to generate ATP, which led us to a hypothesis that HSA cells, neoplastic endothelial cells, highly rely on lactate to regulate transcription.

Here we established canine HSA cell lines and patient-derived xenograft (PDX) models from dog patients and examined the role of histone lactylation under glucose-deprived conditions. To minimize the effect of prolonged cell culture, mechanistic studies were performed with early-passage cells (fewer than 16 passages). Glucose restriction reduced global histone lactylation levels, while lactylation-peaks were redistributed to transcription-start sites (TSSs) of ATF4 regulated stress-response, asparagine synthesis, and immune-related genes. TSSs of stress-response genes were co-occupied with RNA polymerase II phosphorylated at serin 5 (RNAPII-Ser 5) and showed increased transcription, indicating that these lactylation marks activated transcription. [U-^13^C]glutamine tracing revealed *de novo* asparagine synthesis from glutamine under glucose-deprived conditions, and asparagine supplementation modestly activated cell proliferation *in vitro*. In clinical HSA tissues, H3K18la signals were heterogenous, but tumor regions with low-H3K18la signals accumulated M2-like macrophages. Consistently, HSA cells attracted and differentiated macrophages to the M2-like state, suggesting that HSA tumor cells establish pro-tumor microenvironment in low-lactylation regions. Together, our data show that lysine lactylation, possibly histone lactylation, persists even under glucose-deprived conditions, suggesting that tumor cells use lactate to regulate transcription under nutrient-poor conditions.

## RESULTS

### Establishment of canine hemangiosarcoma cell lines and PDX models

We established two canine HSA cell lines from splenic tumors of two dog patients (HU-HSA-2 and HU-HSA-3; patient details in Table S1). Morphologically, both cell lines were spindle-shaped, although HU-HSA-3 cells were approximately twice as large as HU-HSA-2 cells in nuclear and cellular sizes (Fig. 1A). They expressed endothelial-marker genes and proteins (CD31, vWF, and KDR) (Fig. 1B, 1C, and 1D, Table S2). Gene expression profiles, however, were more similar to those of fibroblasts than those of normal femoral and pulmonary arterial endothelial cells, which might reflect undifferentiated features of neoplastic endothelial cells (Fig. 1E, Table S2). When transplanted subcutaneously into nude mice, both cell lines developed tumors that recapitulate morphological features of HSA such as blood-filled capillaries and proliferation of either spindle (HU-HSA-2) or round to oval (HU-HSA-3) tumor cells (Fig. 1F, 1G). PDX models were also generated from three dog patients (HU-HSA-2, 3, and 1). They retained histological features to the corresponding patient tumors and expressed endothelial markers (CD31 and vWF) (Fig. 1G). Short tandem repeat analysis with a commercial canine genotyping kit confirmed the canine origin and unique identity of each cell line and PDX models (Fig. 1H). We used these resources for the following experiments.

**Fig. 1.**
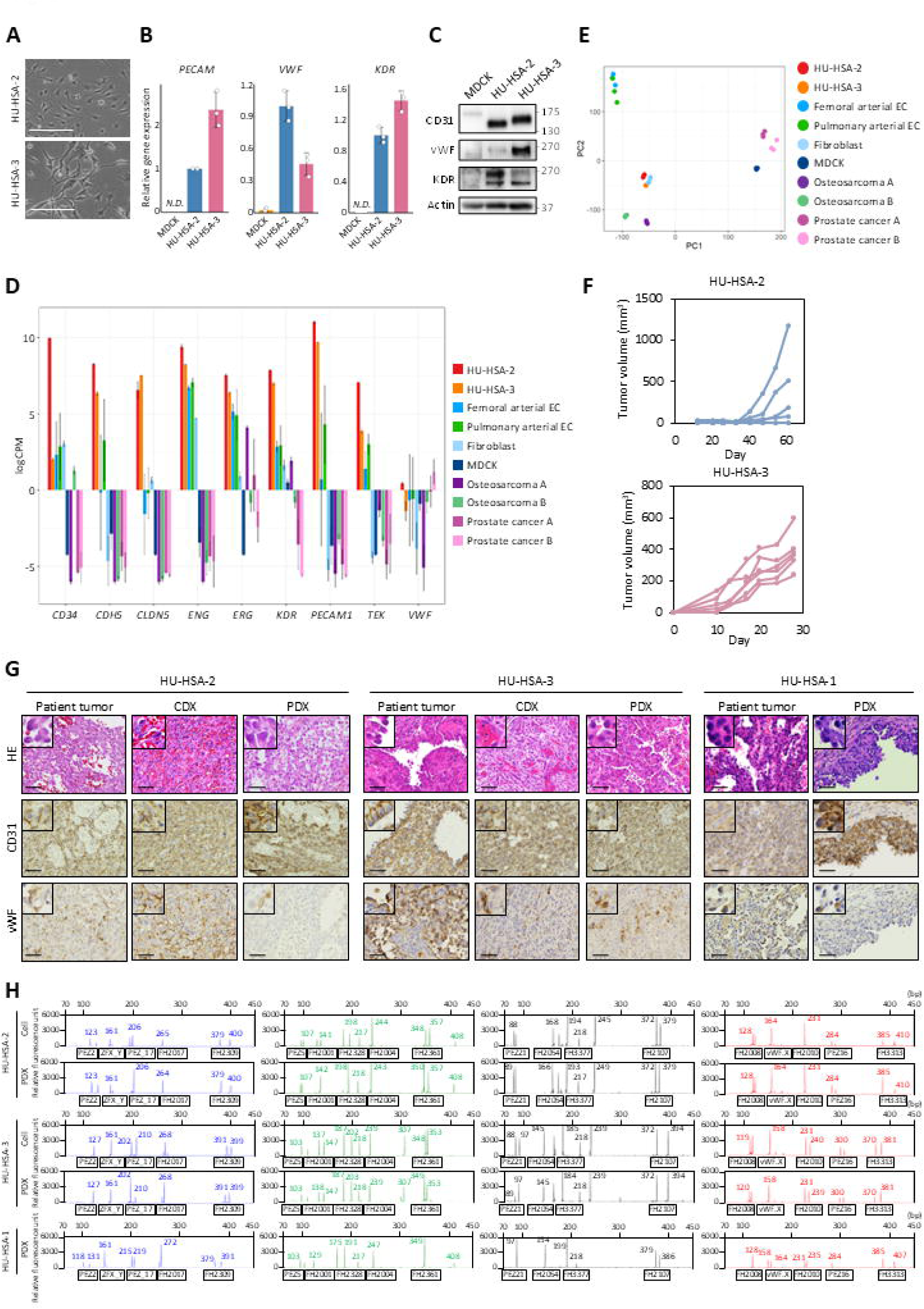
Established HSA cell lines and PDX models retained HSA features. **(A)** Phase contrast images of established HSA cell lines. Scale bars, 250 μm. **(B)** Relative expression levels of endothelial marker genes in HU-HSA-2 and HU-HSA-3 cells with MDCK cells as the negative control. **(C)** Western blot analysis of endothelial marker proteins in HU-HSA-2 and HU-HSA-3 cells. MDCK cells were used as a negative control. **(D)** Principal component analysis of RNA-seq data in HSA cell lines and canine tumor or healthy cells. **(E)** mRNA expression levels of endothelial marker genes in HSA cell lines and canine tumor or healthy cells. **(F)** Tumor growth curves of HU-HSA-2 and HU-HSA-3 cells transplanted into nude mice. Each line represents the volume of tumors formed in individual injection sites (*n* = 6 per group). **(G)** Representative images of HE staining and IHC for the endothelial markers. Images compare the original patient tumor, cell line-derived xenograft (CDX), and PDX for both HU-HSA-2 and HU-HSA-3, and the patient tumor and PDX for HU-HSA-1. Scale bars, 50 µm. **(H)** STR profiles of the established cell lines and their corresponding PDX models. Data are presented as average ± SD from three technical replicates. *N.D.*, Not Detected.

### Glucose is the major source of histone lactylation in HSA cells

To investigate the role of histone lactylation, we first evaluated global histone lactylation levels in the newly established HSA cell lines under regular or nutrient-deficient conditions.

Under the regular culture condition (25 mM glucose, 4 mM glutamine), HU-HSA-2 cells showed a modest increase in global histone lactylation levels compared to normal canine aortic endothelial cells (CnAOEC), while HU-HSA-3 cells exhibited markedly stronger signals (Fig. 2A). Glucose deprivation significantly decreased global pan-H3 and H4 lactylation, H3K18la, and H4K5la levels in both cell lines without altering global histone acetylation levels (H3 acetylation; H3Ac, H4 acetylation; H4Ac) (Fig. 2B), and it resulted in modest cell growth retardation within 96 hours (Fig. 2C). Polyclonal knockout of GLUT1, a glucose transporter, by the CRISPR/Cas9 system also exhibited a significant reduction in global histone lactylation levels (Fig. S1A), although no growth inhibition was observed *in vitro* and *in vivo* (Fig. S1B and S1C). This could be explained by metabolic adaptation acquired during the prolonged establishment period of knockdown cells. In contrast to glucose, glutamine deprivation did not affect global histone lactylation levels (Fig. S1D). These results suggest that glucose is the major source of histone lactylation in canine HSA cell lines.

**Fig. 2.**
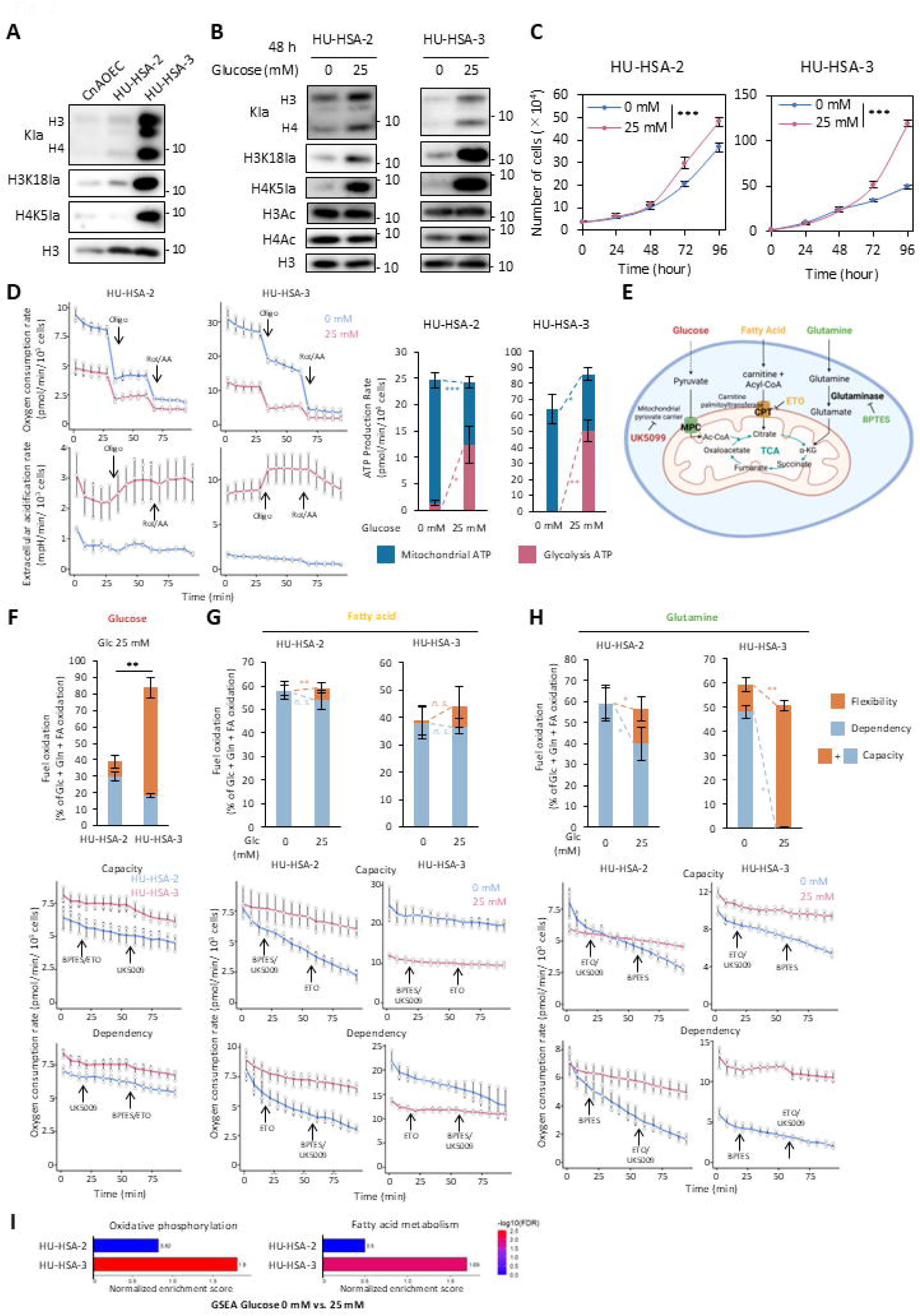
Glucose starvation reduces global histone lactylation levels and reprograms HSA cell metabolism. **(A)** Western blot analysis of histone lactylation and total histone H3 levels in CnAOEC and HSA cell lines. **(B)** Western blot analysis of histone lactylation and acetylation, and total histone H3 levels in HSA cell lines. **(C)** Growth curves of HSA cell lines cultured with or without glucose over 96 hours *in vitro*. **(D)** Extracellular flux analysis of HSA cell lines. (Left) Oxygen consumption rate and extracellular acidification rate traces. (Right) Calculated ATP production rates from mitochondria and glycolysis. **(E)** A schematic diagram of the Seahorse Mito Fuel Flex Test. Created with BioRender. **(F-H)** Mitochondrial fuel oxidation analysis in HSA cell lines. (Upper) The calculated fuel oxidation, dependency, and flexibility for glucose (F), fatty acids (G), and glutamine (G). (Bottom) OCR traces during the sequential inhibition of fuel pathways. **(I)** GSEA from mRNA-seq in HSA cell lines. HSA cells were cultured for 48 hours with or without glucose in (B), (C), (F-I). Data are presented as average ± SD (*n* = 3). *N.s*., not significant. \**P* < 0.05. \*\**P* < 0.01. \*\*\**P* < 0.001. Two-way ANOVA for (C). Student’s *t* test for extracellular flux analyses.

Next, we assessed the metabolic status of HSA cell lines and the effects of glucose deprivation by extracellular flux analyses. Although our HSA cell lines retained endothelial characteristics, they relied comparably on glycolysis and OXPHOS for ATP production (Fig. 2D). Glucose deprivation induced a near-complete metabolic shift toward OXPHOS, while overall ATP production rates remained comparably or slightly decreased (Fig. 2D). To further evaluate nutrient dependency, flexibility, and capacity, we sequentially inhibited major mitochondrial fuel pathways (Fig. 2E). This allowed us to measure cellular reliance on three key mitochondrial fuels—glucose (pyruvate), glutamine, and fatty acids—using specific inhibitors: UK5099 for the mitochondrial pyruvate carrier, BPTES for glutaminase, and Etomoxir (ETO) for carnitine palmitoyltransferase. HU-HSA-3 cells consumed more glucose than HU-HSA-2 (Fig. 2F), which may explain why the higher global histone lactylation levels were observed in HU-HSA-3 (Fig. 2A). Upon glucose deprivation, HSA cell lines became increasingly dependent on fatty acid and glutamine oxidation (Fig. 2G and 2H), probably to compensate for the loss of glucose input. Consistent with these results, gene set enrichment analysis (GSEA) from mRNA-seq showed that HU-HSA-3 cells activated genes related to OXPHOS and fatty acid oxidation, although similar changes were not observed in HU-HSA-2 cells (Fig. 2I). This discrepancy might be associated with more robust alterations of nutrient-dependency in HU-HSA-3.

Taken together, our results demonstrate that HSA cells exhibit elevated global histone lactylation levels, that glucose is the major source of this modification, and that glucose restriction shifted cellular metabolism toward mitochondrial respiration by becoming more dependent on fatty acid and glutamine.

### Lysine lactylation is enriched at TSSs and modulates gene expression during glucose starvation

To determine whether robust lactate reduction by glucose deprivation affects transcriptional regulation, we performed CUT&Tag assay using antibodies against lysine lactylation (Kla), H3K4me3, and H3K27ac in HU-HSA-2 cells cultured for 48 hours with or without glucose. Glucose starvation increased Kla signals at promoter regions (≤ 1 kb), while distributions of H3K4me3 and H3K27ac were not changed (Fig. S2A). Further analysis including HU-HSA-3 confirmed that Kla signals were strongly enriched around TSSs in both HU-HSA-2 and HU-HSA-3 cells under glucose-deprived conditions (Fig. 3A, S2B and S2C). In contrast, H3K4me3 and H3K27ac did not exhibit such localized enrichment with exception for H3K4me3 enrichment at TSSs in HU-HSA-3 (Fig. 3A, S2B, S2C and S3A). We then analyzed co-occupancy of Kla, H3K4me3, and RNAPII-Ser5 around TSSs to assess the transcriptional competence of Kla signals on TSSs in HU-HSA-3 cells under glucose-deprived conditions.

**Fig. 3.**
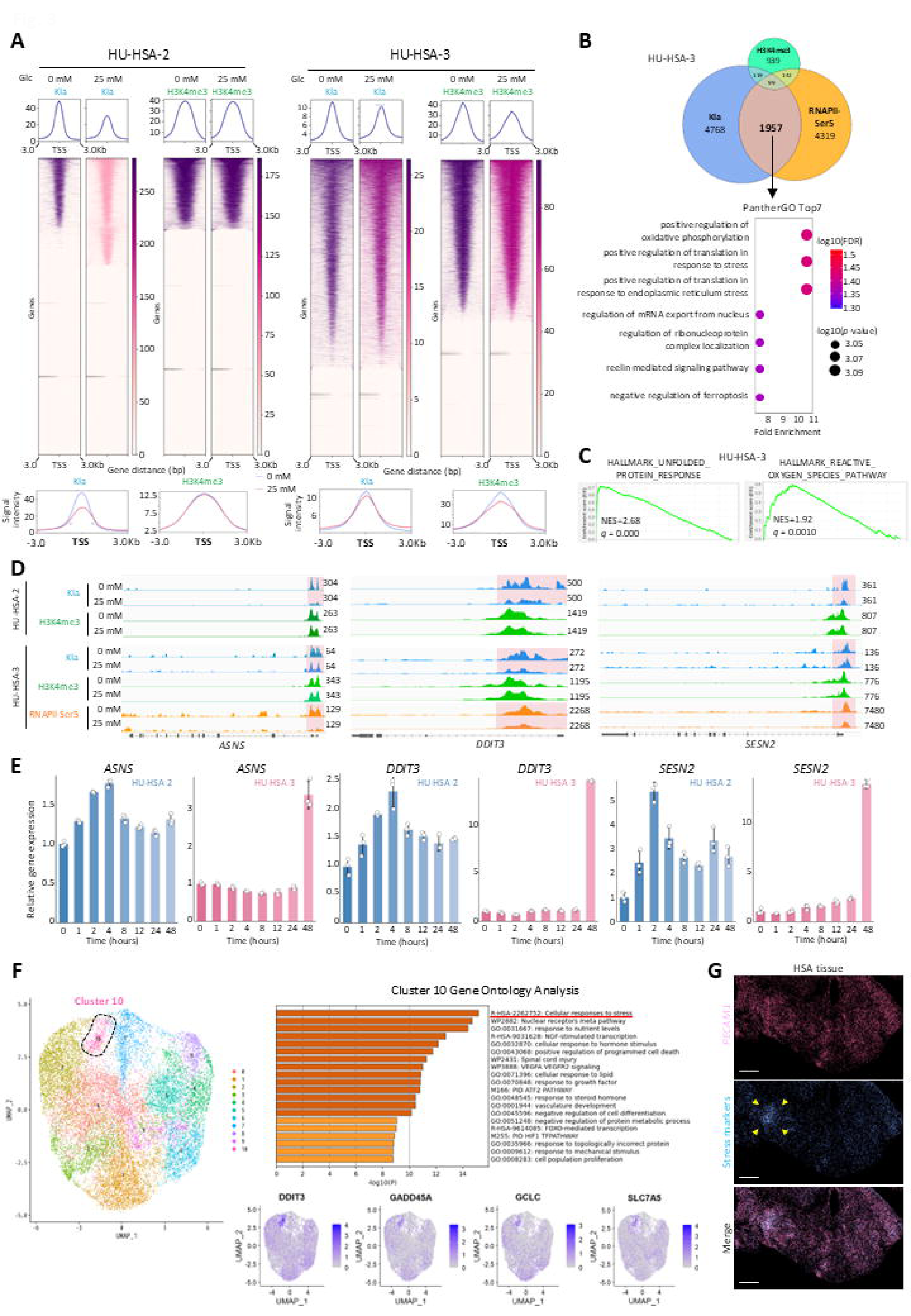
Kla is enriched at TSSs of stress-response genes and promotes their transcription under glucose starvation. **(A)** Composite profile plots (upper), heatmaps (middle) and merged profile plots (lower) around TSSs. **(B)** (Upper) Venn diagram illustrating the overlap of genes with enriched Kla, H3K4me3, or RNAPII-Ser5 at TSSs in HU-HSA-3 cells under glucose starvation. (Lower) Panther Gene Ontology analysis of the 1,857 genes co-enriched for Kla and RNAPII-Ser5 but not H3K4me3. **(C)** GSEA plots of mRNA-seq in glucose-starved HU-HSA-3 cells. NES, Normalized Enrichment Score. **(D)** Genome tracks showing CUT&Tag signal at representative stress response gene loci in HSA cell lines cultured with or without glucose. **(E)** Time-course analysis of relative expression levels of stress response genes in HSA cell lines following glucose starvation. **(F)** Integrated single-cell transcriptomic analysis of two HSA-PDX tumors. (Left) UMAP plot of all tumor cells. (Upper-right) Metascape analysis for cluster 10. (Lower-right) UMAP plots of representative stress-response genes. **(G)** Spatial transcriptomic analysis of an HSA patient tumor tissue. Visualization of *PECAM1* (a marker for HSA tumor cells), stress marker gene expressions, and the merged image. Arrow heads indicate a stress response cluster. Data are presented as average ± SD (*n* = 3).

Interestingly, the largest overlap (1,957 genes) was detected between Kla and RNAPII-Ser5P, whereas only 142 genes were overlapped for H3K4me3 and RNAPII-Ser5P and 99 genes for all three marks, suggesting that Kla activates gene expressions independent from H3K4me3 (Fig 3B, upper). PANTHER Gene Ontology analysis of the genes overlapped with Kla and RNAPII- Ser5 signals enriched gene sets associated with positive regulation of OXPHOS and stress responses (Fig. 3B, lower). Consistently, GSEA on mRNA-seq data showed significant enrichment of OXPHOS and stress-response signatures under glucose starvation (Fig. 2I and 3C). Further examination of representative stress-response genes co-enriched with Kla and RNAPII-Ser5 at their TSSs; *ASNS* (amino-acid deprivation), *DDIT3* (endoplasmic reticulum stress), and *SESN2* (oxidative stress), confirmed concurrent enrichment under glucose-deprived conditions (Fig. 3D). Expression of these and other stress-response genes were upregulated in HU-HSA-3 cells 48 hours after starting glucose starvation (Fig. 3E and S3B). HU-HSA-2 cells also increased these genes expressions, while transcriptional activations peaked 4 hours after starting glucose deprivation (Fig. 3E and S3B). Kla enrichment at TSSs was already observed at 4 hours after starting glucose starvation in HU-HSA-2 cells (Fig. S3C), suggesting that this cell line responded to glucose-deprivation faster than HU-HSA-3 cells.

Next, to evaluate whether similar responses occur *in vivo*, we performed single-cell RNA-seq (scRNA-seq) on two HSA PDX tumors (HU-HSA-1 and HU-HSA-3) and spatial transcriptomic on an HSA patient tumor. The scRNA-seq analysis identified a tumor cell population featured with stress-responses genes including those enriched with Kla and RNAPII- Ser5 signals on their TSSs (Fig. 3F). Of the 227 genes defining cluster 10, 44 were concurrently enriched with Kla and RNAPII-Ser5 at their TSSs (Fig. S3D). We then mapped tumor cells that expressed stress-response genes from the overlap (*ATF3, DDIT3, DDIT4, GADD45A,* and *GCLC*) as well as *ATF4* and *ASNS,* and found that they formed cluster-like regions rather than randomly scattered, suggesting that their distribution is shaped by the local microenvironment (Fig. 3G). These findings indicate that these stress response pathways are also active *in vivo*.

Collectively, our data indicates that glucose starvation leads to Kla enrichment at TSSs of OXPHOS and stress-response genes in HSA, and that this enrichment correlates with increased transcription.

### ATF4 and acute glucose removal are required to induce stress responses

To explore upstream regulation, we focused on ATF4, one of the master regulators of stress-response genes. CUT&Tag revealed robust Kla enrichment at the TSSs of ATF4 48 hours after glucose withdrawal, whereas RNAPII-Ser5 enrichment was decreased (Fig. 4A). Polyclonal ATF4 knockout in HU-HSA-3 cells significantly dampened the upregulation of ATF4-depndent stress response genes induced by glucose deprivation, but it did not affect ATF4-independent stress-response genes such as *NQO1* and *GCLC* (Fig. 4B and 4C). ATF4 loss did not affect short-term cell proliferation under either glucose-starved or normal culture conditions (Fig. 4D), indicating that ATF4-regulated stress responses were negligible for proliferation or other stress-response regulators compensated for ATF4 loss. In the time course analysis of glucose deprivation, global histone lactylation levels started decreasing after 24 hours in HU-HSA-2 cells and 8 hours in HU-HSA-3 cells (Fig. 4E). Although RNAPII-Ser5 was not enriched on TSSs of ATF4, its protein level was increased in HU-HSA-2 cells within 1 hour and in HU-HSA-3 cells by 48 hours after glucose removal with subsequent expression of ASNS (Fig. 4E). Stepwise glucose dilutions indicated that ATF4 expression was induced at higher glucose concentrations in HU-HSA-3 than HU-HSA-2 cells, although histone lactylation levels were reduced in both cell lines even at 2.5 mM glucose (10 times dilution, Fig. 5A). *ATF4* and *ASNS* expressions showed inverse correlation with glucose concentrations in both cell lines; however, the extent of upregulation was greater in HU-HSA-3 cells (Fig. 5B). These results suggest that HU-HSA-2 and HU-HSA-3 cell lines have different sensitivities to glucose deprivation, which could reflect their different dependency on glucose for ATP production (Fig. 2D).

**Fig. 4.**
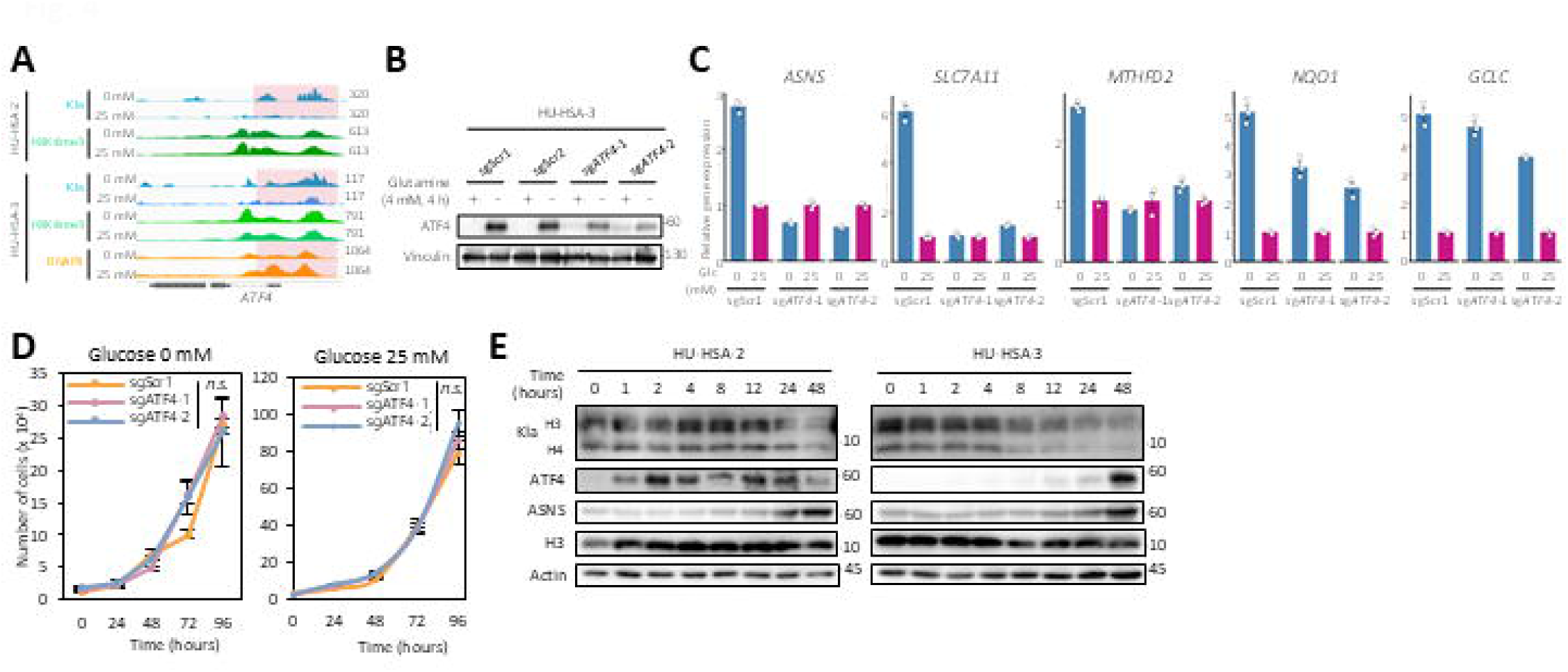
ATF4 is required to induce stress responses under glucose starvation. **(A)** Genome tracks showing CUT&Tag signal at *ATF4* in HSA cell lines cultured for 48 hours with or without glucose. **(B)** Western blot analysis of Kla, ATF4, ASNS, total histone H3, and Actin over a 48-hour time course of glucose starvation in HSA cell lines. **(C)** Western blot analysis of ATF4 in ATF4 polyclonal knockout HU-HSA-3 cells and the scramble controls cultured without glutamine for 4 hours to induce ATF4 expression. **(D)** Relative mRNA expression of key stress response genes in scramble control and sgATF4 expressing HU-HSA-3 cells cultured for 48 hours with or without glucose. Gene expression was normalized to the scramble control group under each respective glucose condition. **(E)** *In vitro* growth curves of scramble control and sgATF4 expressing HU-HSA-3 cells cultured over 96 hours with or without glucose. Data are presented as average ± SD (*n* = 3). *N.s*., not significant, two-way ANOVA.

**Fig. 5.**
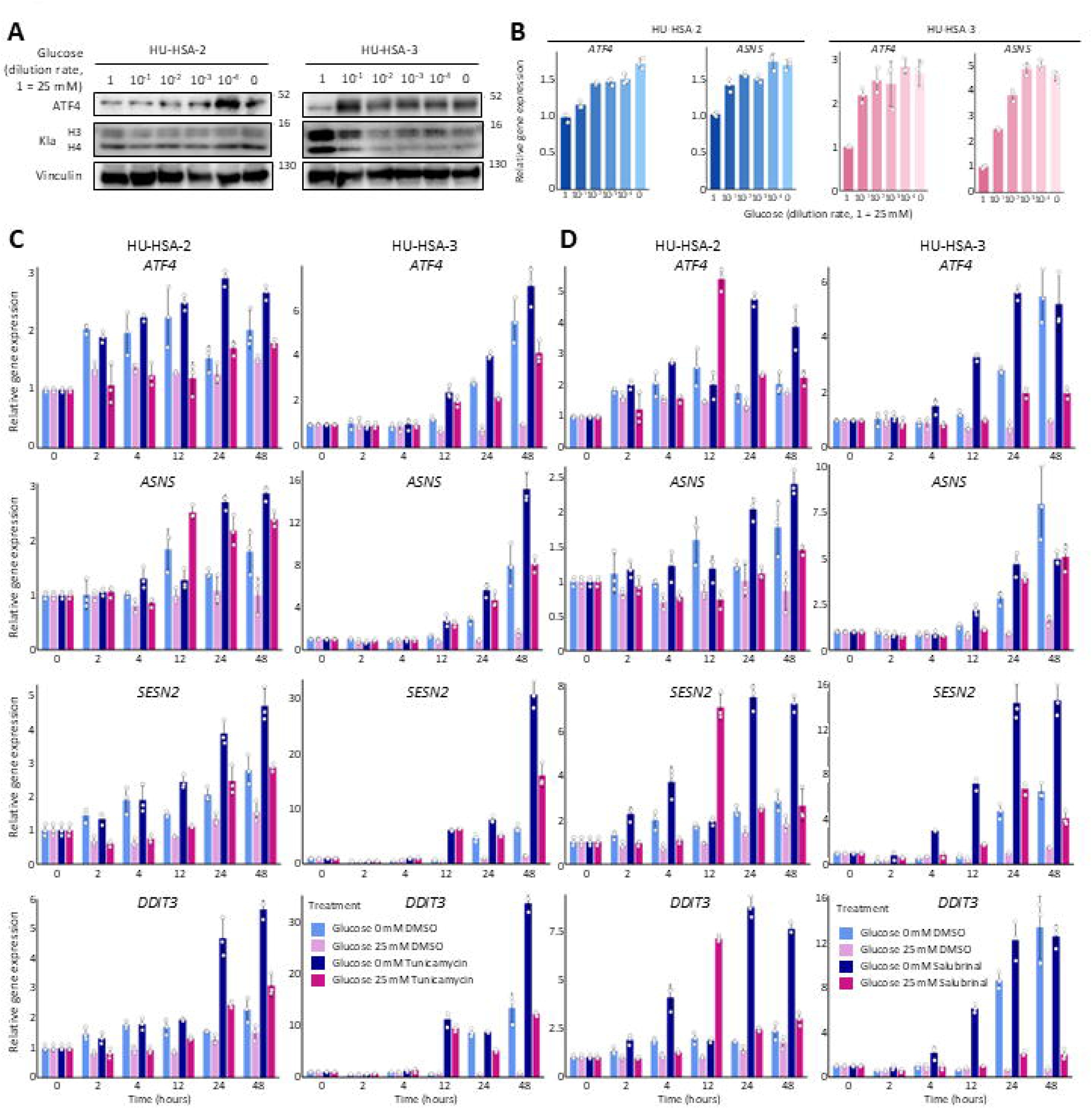
Acute glucose withdrawal is required for inducing stress responses in HSA cells. **(A)** Western blot analysis of ATF4, Kla, and Vinculin in HU-HSA-2 and HU-HSA-3 cells. Cells were cultured for 4 hours (HU-HSA-2) or 48 hours (HU-HSA-3) in medium with serially diluted glucose concentrations from 25 mM to 0 mM. **(B)** Relative mRNA expression levels of *ATF4* and *ASNS* in HU-HSA-2 and HU-HSA-3 cells cultured under the same conditions as in (A). **(C, D)** Time-course analysis of the relative mRNA expression of *ATF4*, *ASNS*, *SESN2*, and *DDIT3* in HU-HSA-2 and HU-HSA-3 cells. Cells were cultured in medium with or without glucose and treated with either a vehicle control (DMSO), 0.06 µg/mL Tunicamycin (C), or 10 µM Salubrinal (D). Data are presented as average ± SD (*n* = 3).

We also limited glucose uptake by polyclonal knockout of GLUT1 (*SLC2A1*) and by treating cells with a GLUT1 inhibitor BAY876. Both treatments significantly reduced global histone lactylation levels (Fig. S1A and S4A), yet they failed to trigger stress-response gene expressions (Fig. S4B and S4C). Although BAY876 treatments slightly increased their expressions (< two-fold) in HU-HSA-2 cells, GLUT1 inhibition could be sufficient to reduce global histone lactylation levels but not to induce stress responses. As we described above, long-term culture during knockout cell-line establishment might have allowed metabolic adaptation. BAY876 treatment could provide partial inhibition, either because of insufficient inhibition of GLUT1 or compensation by other glucose transporters such as GLUT3 or GLUT4. Thus, acute and substantial glucose removal appears necessary to induce stress responses.

Next, to test whether acute glucose starvation is required for inducing stress responses, we treated the cells with tunicamycin and salubrinal. Tunicamycin directly induces ER stress by inhibiting N-linked glycosylation (*31*), whereas salubrinal prolongs stress responses by inhibiting eIF2α dephosphorylation, thereby increasing ATF4 protein levels (*32*). We used these compounds at a low-dose (IC_25_) to mimic glucose-deficient cultures we tested, which induced only a slight cell proliferation delay (Fig. S4D). These treatments activated *ATF4, ASNS, SESN2,* and *DDIT3* expressions under regular culture conditions, confirming stress responses were induced (Fig. 5C, D, dark red). Glucose deprivation alone induced almost comparable expression levels to those in tunicamycin or salubrinal treatments, whereas modest additive effect was observed with drug treatments at 48 hours post-treatment (Fig. 5C and 5D). In addition, these treatments induced gene expressions slightly earlier than DMSO controls in HU-HSA-3 cells, yet expression levels eventually reached the same levels by 48 hours post-treatment. These results suggest that acute glucose deprivation alone is sufficient to induce robust ATF4-regulated stress responses.

Taken together, glucose-starvation-induced stress responses are mostly ATF4 dependent and require acute and substantial glucose withdrawal, while global histone lactylation loss alone is insufficient to trigger them.

### HSA cells activate *de novo* asparagine synthesis from glutamine to survive in glucose-deprived conditions

So far, we have shown that glucose deprivation activates transcription of ATF4-mediated stress-response genes including the asparagine synthetase, ASNS. CUT&Tag analysis also revealed Kla enrichment at the TSSs of genes involved in asparagine and aspartate biosynthesis under glucose-deprived conditions (Fig. 6A). We therefore hypothesized that *de novo* asparagine synthesis contributed to metabolic adaptation during acute glucose starvation. Given that glucose withdrawal increased OXPHOS activity and glutamine dependence in HSA cells (Fig. 2D, 2H), glutamine could be used for asparagine synthesis via anaplerosis (Fig. 6B).

**Fig. 6.**
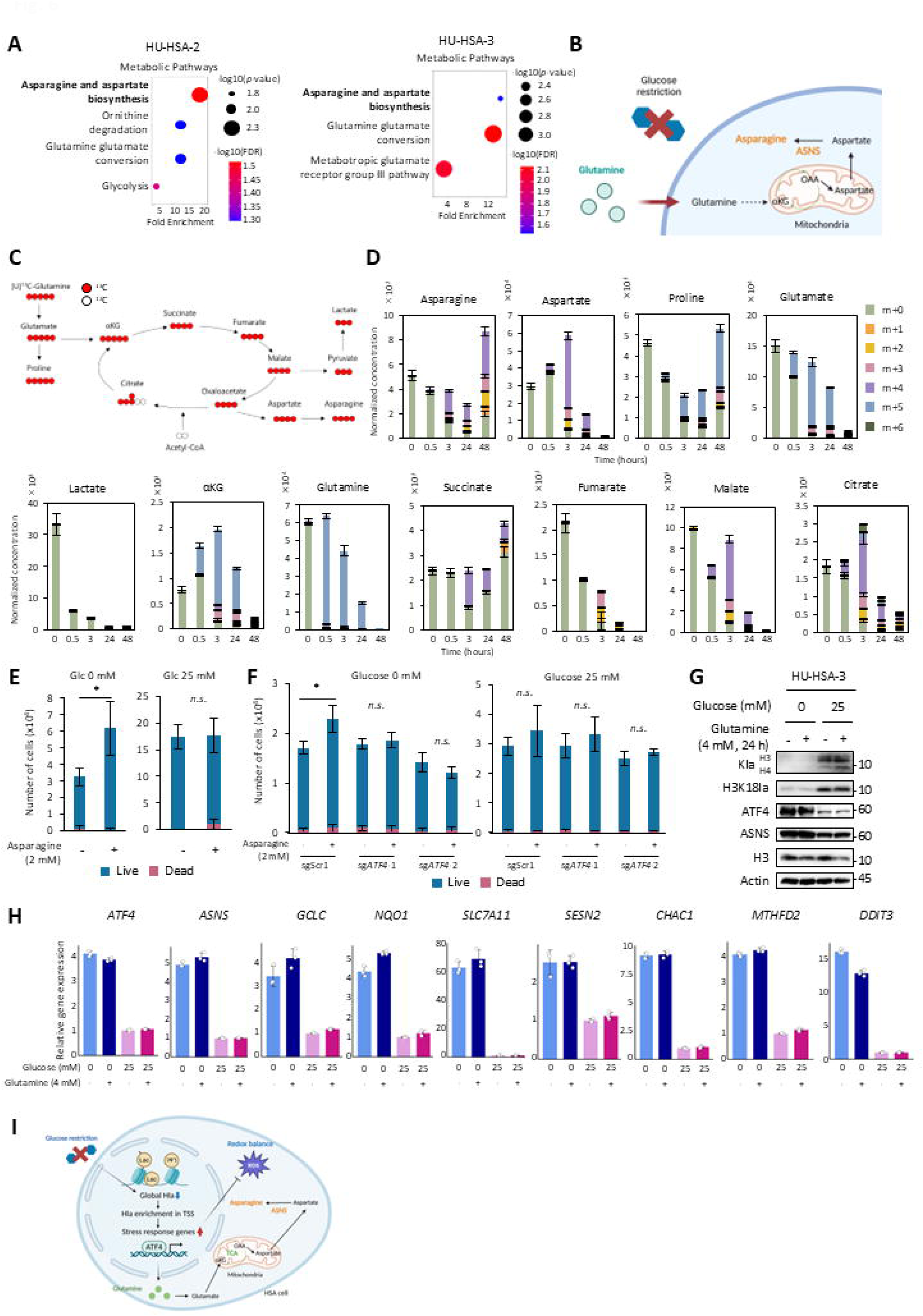
HSA cells utilize glutamine-derived asparagine for survival under glucose starvation. **(A)** Panther Pathway analysis of genes with Kla enrichment at the TSSs in glucose-starved HSA cells. **(B)** Diagram of *de novo* asparagine synthesis via the TCA cycle under glucose restriction. Created with BioRender. **(C)** Schematic diagram of the [U-^13^C]Glutamine tracing. **(D)** Normalized concentrations of metabolites in HU-HSA-3 cells cultured under glucose starvation switching to medium containing [U-^13^C]Glutamine. **(E)** Cell proliferation assay of HU-HSA-3 cells cultured for 72 hours with or without glucose and supplemented with or without 2 mM asparagine in 1% FBS. **(F)** Cell proliferation assay of scramble control and sgATF4 expressing HU-HSA-3 cells cultured under the same conditions as (E). **(G)** Western blot analysis for Kla, H3K18la, ATF4, and ASNS in HU-HSA-3 cells treated as in (F). **(H)** Relative expression levels of stress response genes in HU-HSA-3 cells cultured for 48 hours with or without glucose. 4 mM glutamine was supplemented for the final 24 hours of culture. **(I)** A proposed model summarizing the adaptive response of HSA cells to glucose starvation. Created with BioRender. Data are presented as average ± SD (*n* = 3). \**P* < 0.05; *n.s.*, not significant. Student’s *t* test.

To trace glutamine-derived carbon, we conducted isotope-tracing metabolomic analysis using [U-^13^C]glutamine for 48 hours in HU-HSA-3 cells (Fig. 6C). The results confirmed that glutamine was used to fuel the TCA cycles since ^13^C label appeared in TCA cycle intermediates within 0.5 hours (Fig. 6D). ^13^C incorporation into asparagine started at 3 hours after glucose starvation, and the asparagine concentration significantly increased by 48 hours (Fig. 6D). By contrast, ^13^C enrichment and concentration of aspartate peaked at 3 hours and then decreased gradually, suggesting that aspartate was converted to asparagine by ASNS. Lactate concentration was markedly reduced 0.5 hours after glucose starvation (Fig. 6D). This confirms that Kla enrichment on TSSs was established under low-lactate conditions. Next, we added asparagine (2 mM at a final concentration) to the culture medium under glucose-starved and low-FBS conditions. In this experiment, we reduced FBS concentrations to minimize the effect of asparagine supplied from serum and to examine the direct effect of asparagine supplementation. Asparagine supplementation modestly increased HSA cell proliferation rates under glucose-deprived conditions (Fig. 6E and S4A), and its effect was abolished by ATF4 reduction (Fig. 6F). In contrast, asparagine supplementation had no impact under normal glucose conditions (Fig. 6E and S5A). Although proline showed a similar trend to asparagine (Fig. 6D), proline supplementation did not accelerate proliferation, nor were proline synthesis genes upregulated (Fig. S5B and S5C). These results suggest that glutamine-derived asparagine, produced through the ATF4-ASNS axis, supports HSA cell survival in glucose-deprived conditions.

^13^C tracing also indicated that intracellular glutamine was significantly decreased at 24 hours and nearly depleted at 48 hours (Fig. 6D), which raised a possibility that glutamine deficiency, rather than glucose starvation, induced stress responses. To address this possibility, we added glutamine at 24 hours post-initiation of glucose starvation and subsequently examined stress-response gene and protein expressions. Glutamine supplementation did not affect gene expression changes induced by glucose starvation, global histone lactylation levels, and ATF4/ASNS protein expression levels (Fig. 6G and 6H). These results suggest that these responses are driven primarily by glucose withdrawal not by secondary glutamine depletion during the extended culture period.

Collectively, we demonstrated that HSA cells activate *de novo* asparagine synthesis from glutamine to adapt to glucose starvation, and that ATF4-mediated stress responses and global histone lactylation loss are induced by glucose deprivation itself (Fig. 6I).

### M2**–**like macrophages accumulate around HSA cells with low histone lactylation levels

Finally, we examined histone lactylation levels and its functional implications in patient tumors. Formalin fixed paraffin-embedded blocks of 13 canine splenic HSA cases archived in our laboratory were used for this purpose (Table S1). Immunohistochemistry (IHC) for H3K18la indicated that HSA tumor cells exhibited significantly higher average H3K18la intensities compared to normal endothelial cells (ECs) in 9 out of 13 cases (Fig. 7A, B). In contrast, no such trend was observed in nuclear Kla signals likely because this antibody recognizes broader targets including non-histone proteins (Fig. S6A and S6B). By careful microscopic observation, we recognized heterogeneous H3K18la signal patterns in tumor cells within the same tissues. We then classified tumor cells into low, middle, and high groups based on nuclear H3K18la mean values and visualized their spatial distribution. The results indicated that tumor cells formed clusters with cells of the same group, suggesting that H3K18la signals are associated with spatial factors such as tumor microenvironment (Fig. 7C).

**Fig. 7.**
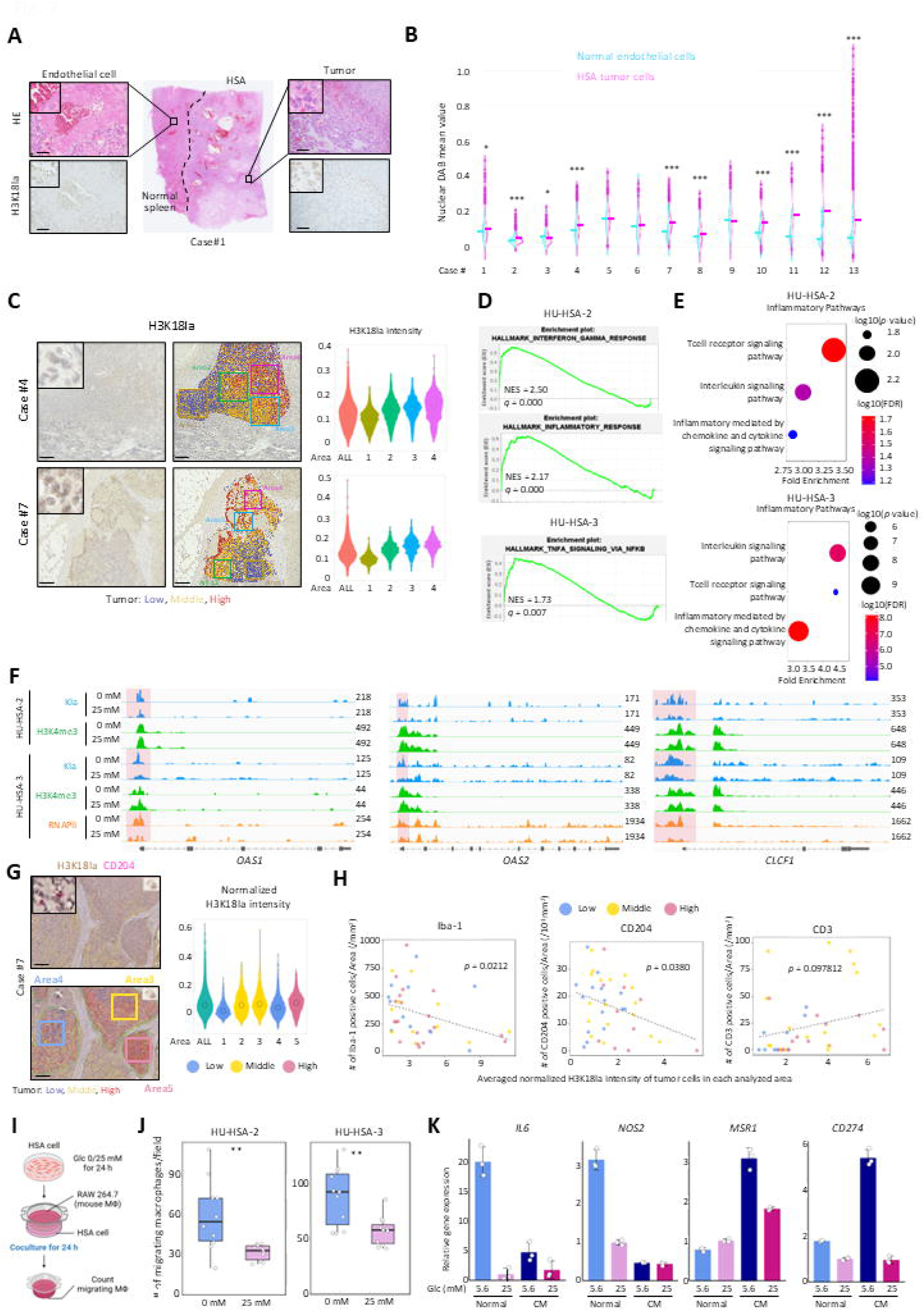
Histone lactylation exhibits heterogeneous distribution, and M2-like macrophages accumulate in low-histone lactylation areas. (A) Representative images of hematoxylin & eosin and H3K18la IHC of HSA and adjacent normal tissues. Scale bars, 100 µm. (B) Quantitative analysis of nuclear H3K18la intensity in HSA and normal endothelial cells. (C) (Left) Representative images of intratumoral H3K18la heterogeneity in HSA tissue. Scale bars, 250 µm. (Right) Violin plots showing H3K18la intensity in total and subregions. (D) GSEA plots from mRNA-seq showing enrichment of inflammatory pathways in glucose-starved HSA cells. (E) Panther Pathway analysis of Kla enriched genes from glucose-starved HSA cells. (F) IGV snapshots of CUT&Tag signals at representative inflammatory gene loci. (G) Representative dual-IHC images for H3K18la and CD204 in HSA tissue. Scale bars, 200 µm. (H) Correlation plots between H3K18la intensity of tumor cells and the number of infiltrating immune cells within each area. (I) Schematic of the migration assay. (J) Box plots of migrating RAW264 cells. (K) Relative mRNA expression in RAW264 cells treated with conditioned medium from HU-HSA-3 cells with or without glucose. Data are presented as average ± SD (*n* = 3). \**P* < 0.05, \*\**P* < 0.01, \*\*\**P* < 0.001. Student’s *t* test.

mRNA-seq and CUT&Tag experiments on glucose-deprived HSA cell lines revealed transcriptional activation and Kla enrichment at the TSSs of inflammation-associated genes (Fig. 7D, 7E and 7F). Based on these findings, we double-stained H3K18la and immune cell markers (Iba-1 for macrophages, CD204 for M2-like macrophages, and CD3 for T cells) to evaluate correlations between histone lactylation and immune responses. To take heterogeneous H3K18la patterns in account, we first classified tumor cells as described above and selected five areas that included all three groups for each tissue sample (Fig. 7G). The results indicated statistically significant negative correlation between average normalized H3K18la signals and Iba-1-positive or CD204-positive cells (*p* = 0.0212 and *p* = 0.0380, respectively), whereas no correlation was observed with CD3-positive cells (Fig. 7H). These findings suggest that macrophages, particularly those with an M2-like phenotype, preferentially infiltrate into tumor regions characterized by low histone lactylation levels.

To further explore the functional interaction between glucose-starved HSA cells and macrophages, we co-cultured HSA cells with murine macrophage cell line RAW264 (Fig. 7I). Glucose-starved HSA cells attracted a significantly higher number of RAW264 cells compared to HSA cells cultured under regular glucose conditions (Fig. 7J). Furthermore, conditioned medium from glucose-restricted HSA cells decreased expression of the M1-like markers (*IL-6* and *NOS2*) and increased expression of M2-like markers (*MSR1* and *CD274*) in RAW264 cells (Fig. 7K). These results indicate that glucose-starved HSA cells attracted macrophages and polarized them towards an M2-like phenotype.

Taken together, we demonstrated that histone lactylation was spatially heterogeneous in HSA tissues, and that tumor regions with low histone lactylation levels recruited M2-type macrophages. Considering these findings together with the *in vitro* co-culture experiment results, HSA cells can create pro-tumor microenvironments in glucose-restricted regions in tumor tissues.

## DISCUSSION

In this study, we established and characterized canine HSA cell lines and PDX models by examining their morphology, gene and protein expression, and STR profiles. Long-term *in vitro* cultures can induce genetic drift, clonal selection, and altered signaling pathway activity, thereby causing tumor cells to lose original characteristics (*33, 34*). This makes it difficult to predict the patient responses from results obtained with long-term cultured cells (*35, 36*). To minimize this disadvantage, in our study, all *in vitro* experiments were conducted with early-passage cultures (fewer than p16). We confirmed that they preserved endothelial phenotypes and developed tumors composed of abundant vascular channels, indicating that our cell lines maintain hemangiosarcoma phenotypes at least the morphological level. In addition to early passage cell lines, PDX models are useful tools to predict how patients respond to potential therapeutics.

Generally, PDX models retain morphology and heterogeneity more faithfully than cultured cells because they were grown in a 3D environment with non-tumor components such as stromal and immune cells (*37*). Indeed, our HSA PDX models recapitulated original patient tumor morphology more accurately than early passage cell lines. Although further characterization is needed, our paired patient-derived HSA models could be useful for basic and translational hemangiosarcoma research. Moreover, from a comparative-oncology perspective, our models are highly relevant to human angiosarcoma studies because canine and human (hem)angiosarcoma share morphological, transcriptional, and genetic similarities (*38, 39*). We believe that these resources will accelerate basic studies and the development of novel therapeutics for both canine and human patients.

Our metabolic analysis revealed that HSA cell lines produce ATP through both glycolysis and OXPHOS. This represents a hallmark of tumor metabolism and distinguishes them from that in actively proliferating normal endothelial cells, which rely almost exclusively on glycolysis (*40*). Although we did not perform a direct comparison, glycolytic flux in HSA cell lines likely surpasses that of normal endothelial cells since HSA cell lines showed high levels of glucose-dependent histone lactylation. Such metabolic divergences are worth exploring to develop effective HSA therapeutics while sparing normal endothelial cells. Glucose withdrawal from culture medium resulted in a redistribution of Kla to TSSs of genes associated with stress responses, asparagine synthesis, and immune responses. Stress-response genes exhibited concomitant enrichment of Kla and RNAPII-Ser5 on TSSs and were transcriptionally upregulated, suggesting that Kla at TSSs is associated with positive transcriptional regulation. This is consistent with the previous findings that histone lactylation is enriched at TSSs and actively regulates gene expression in tumor cells and immune cells (*24, 41, 42*), but these studies were conducted under lactate-rich conditions. Our results indicate that HSA cells can utilize a limited lactate pool to regulate selected genes to respond to glucose/lactate shortage conditions. A limitation of our work is that we were unable to determine whether Kla enrichments on TSSs were established on histones. These signals may reflect lactylation of histones, non-histone proteins, or both. Although we tested an H3K18la antibody for CUT&Tag experiments as reported elsewhere, we failed to obtain enough DNA for sequencing. This might be attributed to insufficient H3K18la levels under glucose-deprived conditions, lot-to-lot variability of the antibody, or species differences. Nevertheless, the observed redistributions of Kla under glucose- deprived conditions could indicate that HSA cells utilize a limited amount of lactate to epigenetically regulate gene expression to adapt to low-glucose environments.

Genes associated with asparagine synthesis and immune responses showed Kla peaks on their TSSs but not over-represented amount the Panther GO term for genes with concomitant enrichment of Kla and RNAPII-Ser5. This means that most genes with Kla peaks on their TSSs do not accompany RNAPII-Ser5 peaks, although some genes such as *OAS1*, *OAS2*, and *CLCF1* exhibited co-enrichment of both signals (Fig. 7F). We speculate that these histone lactylation marks prime specific genes for rapid expression upon a second stimulus. Several reports indicate similar mechanisms in innate immunity. M1 macrophages exposed to bacteria primed wound-healing genes that were later transcribed during M2 polarization (*24*) and trained monocytes/macrophages retain H3K18la as an epigenetic memory that accelerates gene expression upon a secondary stimulus. While our experiments demonstrated that the asparagine pathway and immune responses proved functionally important in HSA cells, we need to determine whether Kla actually primes these genes for rapid activation upon a secondary stimulus by evaluating gene and protein expression time courses. In addition, our CUT&Tag experiment revealed that ATF4 was one of the genes with Kla enrichment at TSSs without RNAPII-Ser5 enrichment. Given that ATF4 protein expression is regulated by translational inhibition (*43*), concomitant enrichment of Kla and RNAPII-Ser5 may not necessarily be important for immediate induction. However, it is possible that Kla at TSSs functions as an epigenetic memory to rapidly supply additional ATF4 protein if ATF4 protein becomes limiting or additional cellular stresses occur. Overall, although further research is required, Kla, possibly histone lactylation, could maintain selected genes in a poised state for future stimuli in tumor cells as well.

In addition to the limitations noted above, our study has several further caveats. First, all metabolic analyses were performed in 2D cell cultures. Tumor cell metabolism *in vivo* is modulated by microenvironmental factors, which results in metabolic dynamics distinct from those *in vitro*. We should interrogate HSA cell metabolism in cell-line xenografts, PDX models, and patient tissues for more physiologically relevant understandings. Second, we could not perform loss- and gain-of-function experiments for histone lactylation. Writers and erasers for histone have been identified, yet they can also modify histone acetylation (*24, 44*). This means that their knockout, overexpression or pharmacological inhibitions can affect multiple histone modifications, which obscures whether Kla peaks on TSSs drive transcription or merely correlate with transcriptionally active epigenetic state. Third, our experiments were limited to canine hemangiosarcoma. Whether the glucose deprivation-associated enrichment of pan-Kla at TSSs is conserved in human angiosarcoma or across other tumor types remains unknown. If conserved, this lactylation-based gene regulation could be targeted broadly. If not, it may be unique to HSA and best pursued as an HSA-specific target. Nonetheless, our data show that lysine lactylation, possibly histone lactylation, persists even under glucose-deprived conditions, suggesting that tumor cells deploy this epigenetic mark to regulate transcription under nutrient-poor conditions.

## MATERIALS AND METHODS

### Study design

This study aimed to investigate the roles of histone lactylation in response to glucose-deprivation in canine HSA. We established HSA cell lines and PDX models from dog patients and for most experiments. Clinical samples and public mRNA-seq data for canine cell lines or healthy tissues were also used. To assess the roles of histone lactylation under glucose deprivation, we performed mRNA-seq, RT-qPCR, western blotting, histopathological analysis, IHC, CUT&Tag, cell growth assay, extracellular flux analysis, scRNA-seq, spatial transcriptomics, migration assay, conditioned medium assay, metabolome analysis, cell viability assay, animal studies, bioinformatic and statistical analysis. CRISPR/Cas9 and lentivirus transduction system were used to establish polyclonal ATF4 knockout cells. Mice were age-, gender-, and genetic background–matched and randomized to different groups of at least two animals per group before the start of each experiment. Sample size was based on prior knowledge of the intragroup variation of tumor growth. Blinding was not done because the information was essential for the staff to conduct the studies. No data were excluded.

### Reagents, kits, and instruments

All reagents, kits, and instruments used in this study are summarized in Table S3 and S4 along with their company information and catalog numbers.

### Establishment and characterization of canine HSA cell lines and PDX models

Canine HSA cell lines and PDX models were established from fresh hemangiosarcoma tissues obtained from dog patients that underwent splenectomy at Hokkaido University Veterinary Teaching Hospital (HUVTH) with written informed consent from the owners and research ethics approval by HUVTH ethics screening committee (2022–005). All cases were confirmed as hemangiosarcoma by two board-certified veterinary pathologists. Patient information is detailed in Table S1. Tumor tissues were used for cell line establishment and PDX development immediately following surgical resection.

For cell culture, tumor tissues were washed with PBS and then mechanically minced into small fragments (∼1-2 mm cubes) with sterile scalpels and scissors. Tissue fragments were washed with PBS and then treated with an NH₄Cl buffer {8.3% NH4Cl and 170 mM Tris-HCl (pH 7.5)} with gentle agitation twice to remove red blood cells. Following this, tissue fragments were enzymatically digested in Dulbecco’s Modified Eagle Medium (DMEM) containing 3 mg/mL collagenase I for 50 min at 37°C with intermittent mixing. Then, the digested tissues were homogenized by sequential passage through 18G and 23G needles, followed by filtration with a 70 µm cell strainer to remove remaining tissue fragments. The PBS wash and RBC lysis steps were repeated, and then the isolated cells were seeded and maintained in 10 cm culture dishes with DMEM supplemented with 10% Fetal Bovine Serum (FBS) and Penicillin-streptomycin (100 units/mL penicillin, 100 µg/mL streptomycin) (Complete DMEM) at 37°C in a humidified 5% CO₂ incubator. Cells were used for this study after 10 passages by which time the cultured cells appeared homogeneous and proliferated stably. To validate that the established cell lines were hemangiosarcoma cells, endothelial marker gene and protein expressions were assessed by RT-qPCR and western blotting. Subcutaneous transplantation into 6- to 8-week-old female KSN/Slc mice was conducted to evaluate their tumorigenicity and morphological characteristics of cell-derived tumors. Cell line authentication was performed by short tandem repeat (STR) analysis using the Canine Genotypes Panel 2.1 Kit.

For PDX establishment, tumor tissues were fragmented into approximately 2-3 mm cubes and then subcutaneously transplanted into both flanks of 6- to 8-week-old male or female KSN/Slc mice according to the animal study protocol described below. 0.2 mg/kg intraperitoneal injection of meloxicam was done for analgesia on the day of surgery and again the following day. Tumors were resected when the volume reached 1 cm^3^, cut into small pieces again, and then transplanted into other mice. This passage procedure was repeated three times to designate the passaged tissues as canine HSA PDX models. Histopathological examination and immunohistochemistry for endothelial markers were performed to confirm whether HSA PDX models retain patient tumor features. Authentication was done by STR analysis using the Canine Genotypes Panel 2.1 Kit.

### Cell line and cell culture

Canine HSA cell lines, HU-HSA-2 and HU-HSA-3, were established as described above. Madin-Darby Canine Kidney cells (MDCK) were obtained from ATCC. CnAOEC was purchased from Cell Applications. Human embryonic kidney 293T cells and RAW264 cells were obtained from RIKEN Bioresource Center. Normally, cells were cultured with DMEM high glucose supplemented with 10% FBS and penicillin–streptomycin at 37°C with 5% CO_2_. In specific experimental conditions, DMEM without glucose and/or glutamine was used, or FBS concentration was reduced to 1%. Cell culture dishes were coated with 0.1% gelatin when culturing canine HSA cell lines. All cell lines used in this study were confirmed to be free of mycoplasma contamination by PCR (*45*).

### Animal studies

All mouse experiments including PDX model establishment were performed under the guidelines of Hokkaido University (protocol number: 20-0083 and 21-0062), which follow the ARRIVE guidelines. Five-week-old female KSN/Slc mice (Japan SLC, Inc. Shizuoka, Japan) were used for cell line transplantation experiments. Three million HU-HSA-2 cells or one million HU-HSA-3 cells were inoculated subcutaneously in both flanks of KSN/Slc mice anesthetized with 0.3 mg/kg medetomidine, 4 mg/kg midazolam and 5 mg/kg butorphanol. After tumor cell inoculation, mice were awoken by intraperitoneal injection of 3 mg/kg atipamezole. Tumor volumes were calculated using the formula: volume = (length × width^2^)/2. Mice were euthanized with CO_2_ when tumors reached 1 cm^3^ in volume.

### Statistical Analysis

All statistical analyses were performed using R software (version 4.5.0; R Foundation for Statistical Computing, Vienna, Austria). A *p* value of less than 0.05 was considered statistically significant. Data distribution was first assessed for normality using the Shapiro-Wilk test. For comparisons between two independent groups, the two-tailed Student’s *t* test was used for normally distributed data. For comparisons among more than two groups, one-way analysis of variance (ANOVA) was performed, followed by Dunnett’s post-hoc test to compare each experimental group against the control group. For experiments involving two independent variables, two-way ANOVA was used, followed by an appropriate post-hoc test for specific comparisons. *In vivo* tumor growth curves were analyzed using two-way ANOVA with repeated measures, and differences at the final time point were assessed using Dunnett’s multiple comparisons test. Correlations between Hla intensity and immune cell density were assessed using Pearson’s correlation coefficient.

## List of Supplementary Materials

Materials and Methods

Figs. S1 to S5

Tables S1 to S6

References (46–73)

## Acknowledgments

We acknowledge the efforts of Drs. Mitsuyoshi Takiguchi and Hironobu Yasui, Faculty of Veterinary Medicine, Hokkaido University and for giving useful pieces of advice and constructive discussion. We are grateful to all the members of the Laboratory of Comparative Pathology, Faculty of Veterinary Medicine, Hokkaido University for their helpful discussions, encouragement, and support. We sincerely thank all canine patients whose tumor tissues gave rise to the tumor cell lines and PDX models analyzed in this study, and we are deeply grateful to their owners for their generous cooperation and courtesy in supporting this research. We also gratefully acknowledge the following individuals whose contributions to our crowdfunding project helped make this research possible: Akinori Ariji (member of the amateur pro-wrestling group “Nariagari”), Hidetaka Kano, Hirokazu Enomoto, Hisashi Ishihara, Ikuo Konishi, Jun Murayama, Kaoru Miyoda, Kayo Watanabe, Keisuke Okutani, Lemi Shinozaki, Mariko Nishikida, Mayumi Befu, Midori Okutani, Sachiko Hashimoto, Satoru Okita, Teruko Michiduka, Tomoichi Enomoto, Toshiaki Michiduka, Yoji Hibiki, and Yuki Maki, as well as the many other donors who preferred to remain anonymous.

## Funding

This study was supported by the Japan Society for the Promotion of Science (JSPS) KAKENHI Grant-in-Aid for Scientific Research (C) 22K06020 (to K.A.) and Grant-in-Aid for JSPS Research Fellows 23KJ0056 (to T.S.); the Clinical Research Promotion Research Grant, Faculty of Veterinary Medicine, Hokkaido University (to K.A.); a crowdfunding project to advance basic research on canine HSA (Readyfor, https://readyfor.jp/projects/hsa/announcements/238254) (to K.A.); and donations from the general public.

## Author contributions

T.S. and K.A. designed the research. K.A. conceived and supervised the project. T.S. performed most experiments. T.S. and K.A. conducted bioinformatic analyses and animal studies. T.S., K. Heishima, Y.O-O., and K.A. developed the metabolic-analysis methodology. T.S., J.Y., M.Y., P.X., Q.Y., and K.A. established the CUT&Tag protocol and analyzed the data. T.S. and M.S. performed single-cell RNA-seq analysis. R.K., S.K., and K. Hosoya provided clinical samples and patient information. T.S. and K.A. acquired funding. T.S. and K.A. wrote the original draft. All authors reviewed and edited the manuscript.

## Competing interests

Authors declare that they have no competing interests.

## Data and materials availability

The datasets generated and/or analyzed during the current study are available from the corresponding author on reasonable request. RNA-seq and CUT&Tag data were uploaded on Gene Expression Omnibus (GSE304507 and GSE304509). Custom codes used for data analysis are available at Zenodo (DOI 10.5281/zenodo.16813072). All noncommercially available new materials, including constructs, cell lines and PDX models, that Hokkaido University has the right to provide will be made available to nonprofit or academic requesters upon completion of a standard material transfer agreement. Requests for materials may be made by contacting K.A.

## Notes

### Competing Interest Statement

The authors have declared no competing interest.

