## Supplemental information for "Lysine lactylation regulates ATF4-mediated stress responses under glucose starvation in canine hemangiosarcoma"

Tamami Suzuki *et al.*

**The PDF file includes:**

Materials and Methods

Figs. S1 to S6

Tables S1 to S6

References (46-73)

### MATERIALS AND METHODS

#### Western blotting

SDS lysis buffer {2% SDS, 50 mM Tris-HCl (pH6.8), and 1 mM EDTA (pH 8.0)} was added to cultured cells after washing them with ice-cold PBS twice. Cell lysates were then sonicated using BRANSON Sonifier 450 for 2 seconds at power 2. Protein concentrations were measured with TaKaRa BCA Protein Assay Kit before adding 4× sample loading buffer {200 mM Tris-HCl buffer (pH 6.8), 8% SDS, 40% glycerol, 1% bromophenol blue, and 20% 2-mercaptoethanol} and denaturing samples at 98°C for 5 min. 2-5 μg of protein was separated on gradient SDS polyacrylamide gels by electrophoresis and transferred to Immobilon-P transfer membranes. Membranes were blocked with 3% skim milk in Tris-buffered saline with 0.05% Tween 20 (TBST) for 1 hour at room temperature (RT) and incubated with primary antibodies diluted in Can Get Signal Solution 1 overnight at 4°C. Membranes were washed with TBST three times before incubating with the corresponding secondary anti-mouse or anti-rabbit IgG antibodies conjugated with horseradish peroxidase in Can Get Signal Solution 2. Signals were developed with Immobilon Western Chemiluminescent HRP substrate and visualized in Image Quant LAS 4000 mini luminescent image analyzer. Captured data were processed using ImageJ (v1.54p) *(46)*. Antibodies used in this study are listed in Table S3.

#### Hematoxylin and eosin staining, and immunohistochemistry (IHC)

Tumor samples were obtained from patients presented to HUVTH with written informed consent (Table S1). Samples were fixed in 10% neutral buffered formalin, dehydrated through an ethanol series, cleared with xylene and infiltrated with paraffin wax in Tissue-Tek VIP5 Jr. Then, the samples were embedded in paraffin wax and sliced into 2 μm-thick sections. For hematoxylin and eosin staining, tissues were deparaffinized with xylene and placed in 99%, 95%, 90%, 80%, and 70% ethanol for 2 min each in this order. After washing out the ethanol with tap water and distilled water (DW), the tissues were stained with hematoxylin for 1 min and then washed with tap water for 5 min. Then, they were stained with eosin for 1.5 min after being placed in 95% ethanol for 2 min. Remaining eosin was washed with 95% ethanol. Afterwards, the tissues were dehydrated with absolute ethanol and cleared with xylene. Finally, the tissues were mounted with Eukitt and covered with cover glasses for histopathological analysis. For IHC, after deparaffinization, the tissues were washed with phosphate buffered saline (PBS) three times, and then antigens were retrieved in citrate buffer pH6.0 while being heated in a pressure cooker for 9 min. Endogenous peroxidases were inactivated with 0.3% $H_2O_2$ in methanol for 25 min at RT before blocking the tissue sections with 10% normal goat serum for single staining and 5% skim milk in PBS for double staining for 30 min at RT. For single staining, tissues were stained with anti-L-Lactyl-Histone H3 Lys18 (H3K18la) rabbit monoclonal antibody or anti-pan-lactylated lysine (Kla) rabbit monoclonal antibody overnight at 4°C. Afterwards, the tissues were washed with PBS three times and stained with goat anti-rabbit IgG conjugated peroxidase for 30 min at RT. After washing the tissues with PBS three times, they were treated with peroxidase-conjugated streptavidin for 10 min at RT. The slides were washed with PBS three times again, and then signals were developed by reaction with 3.3'-diaminobenzidine. For double staining, sections stained with anti-H3K18la antibody were autoclaved in citrate buffer pH6.0 for 2 min to deactivate the first antibody. They were washed with PBS three times and stained with anti-Iba1, CD3, and CD204 antibodies overnight at 4°C after blocking with 5% skim milk in PBS for 30 min. Subsequently, the tissues were washed with Tris-buffered saline (TBS) three times and stained with goat anti-rabbit IgG or goat anti-mouse IgG conjugated alkaline phosphatase (AP) for 30 min at RT. The slides were washed with TBS three times again, and then signals were developed with New Fuchsin solution. To quantify IHC results, slides were scanned with NanoZoomer 2.0-RS and analyzed with QuPath ver.0.5.1 *(47)*. Areas within 500 μm from the tissue edge were not selected to avoid the edge effect. For intensity comparison of endothelial and HSA cells, five tumor areas per slide in each case were randomly selected. Normal endothelial cells were obtained near the tumor areas in the same section. At least total 1,000 tumor cells and 100 normal endothelial cells were analyzed for each case. For comparison of H3K18la and Kla intensity in HSA and endothelial cells, raw values of nucleus DAB OD mean were applied. For correlation analysis of H3K18la tumor intensity and number of immune cells, nucleus DAB OD mean values in tumor cells were normalized against those in normal endothelial cells, and the normalized values were used for quantitative analyses. Then tumor areas within each case (cases #4–13) were stratified based on H3K18la staining intensity. The thresholds for low, middle, and high Hla intensity were manually established for each case based on the overall staining distribution across the entire tumor area. Subsequently, five distinct regions, each larger than 1 $mm^2$, were selected per case, ensuring that low, middle, and high Hla areas were included in the analysis. A positive staining threshold for the AP signal of each immune marker (Iba-1, CD204, CD3) was then determined manually based on visual inspection of representative positive and negative cells. Cells with a cell AP OD mean exceeding this threshold were classified as positive. The software was used to automatically count the number of positive cells within each area, and the results were expressed as the density of positive cells per square millimeter (cells/$mm^2$).

**Cell growth assay**

For cell growth assays, cells were seeded in triplicate onto 12-well plates at a density of $1.0 \times 10^4$ cells per well for experiments under 10% FBS conditions, or $1.2 \times 10^4$ cells per well for experiments that required FBS restriction. Cells were allowed to attach to dishes overnight in complete DMEM. On the following day (day 0), medium was replaced with the media prepared for each experimental condition, which were DMEM supplemented with or without glucose, asparagine (2 mM), and/or proline (5 mM) as indicated in each section. At each time point (0, 24, 48, 72, and 96 hours), cells were washed with PBS and detached using 100 µL of 0.25% Trypsin-EDTA solution with a 5-minute incubation at 37°C. This reaction was neutralized by adding 1 mL of complete DMEM. The number of live cells was then determined using CellDrop BF with trypan blue staining.

**Plasmid construction and transfection**

Single guide RNAs (sgRNAs) targeting genes of interest were designed using publicly available tools, including the CRISPR gRNA design tool from Horizon Discovery (https://horizondiscovery.com/ja/ordering-and-calculation-tools/crispr-design-tool), CRISPOR (http://crispor.tefor.net/) *(48)*, and CRISPR direct (https://crispr.dbcls.jp/) *(49)*. To minimize off-target effects, sgRNAs with high specificity and efficiency scores were selected. Oligonucleotides for each selected sgRNA were synthesized, annealed, and cloned into the lentiCRISPRv2 plasmid (a gift from Feng Zhang, Addgene plasmid #52961; RRID: Addgene_52961) *(50)* according to the provider's protocol. The oligonucleotide sequences used for generating each knockout construct are listed in Table S6.

Lentiviral particles were produced by transfecting lentivirus plasmids into 293T cells. Briefly, 293T cells were seeded in 6-well plates and grown to approximately 60-70%

confluency. Cells were then co-transfected with a mixture of 4 μg of the lentiCRISPRv2-sgRNA plasmid, 0.5 μg of the packaging plasmid pCAG-HIVgp (RIKEN BRC, cat. RDB04394) *(51)*, and 0.5 μg of the envelope plasmid pCMV-VSV-G (RIKEN BRC, cat. RDB04392), both of which were provided by the RIKEN BRC through the National BioResource Project of the MEXT/AMED, Japan, using Lipofectamine 3000 and P3000 reagent according to the manufacturer's protocol. Forty-eight hours post-transfection, the supernatant containing lentiviral particles was harvested, filtered through a 0.45 μm pore filter, and supplemented with polybrene to a final concentration of 10 μg/mL. For infection, $2 \times 10^4$ HU-HSA-2 or HU-HSA-3 cells were incubated with the virus-containing supernatant for 8 hours. Following transfection, the viral medium was replaced with complete DMEM, and the cells were cultured for 72 hours. Subsequently, cells stably expressing Cas9 and sgRNAs were selected by culture in complete DMEM containing puromycin at a concentration of 2 μg/mL for HU-HSA-2 or 4 μg/mL for HU-HSA-3.

**Extracellular flux analysis**

Extracellular flux experiments were performed according to the manufacturer’s protocol. Briefly, $2.5 \times 10^3$ HU-HSA-2 or $2.0 \times 10^3$ HU-HSA-3 cells were seeded on assay plates coated with 0.1% gelatin and incubated overnight. On the next day, the medium was replaced with complete DMEM with or without glucose. After 48 hours incubation, the medium was replaced with FBS-free DMEM for flux analysis, which contained 4 mM L-glutamine alone or with 25 mM D(+)-glucose. Cells were incubated for at least 1 hour at 37°C without $CO_2$ regulation. Assay solutions were loaded on each assay cartridge well. For ATP rate assay, 1.5 μM Oligomycin and 0.5 μM Rotenone/antimycin A were applied. For mitochondrial oxidation assay,

3 μM BPTES, 2 μM UK5009 and 4 μM Etomoxir were applied. Oxygen consumption rate and extracellular acidification rate were measured using Seahorse XFp Analyzer. For normalization, cells were stained using Hoechst 33342 after measurement, and the well images were acquired using EVOS FL at 10× magnification. For each well, three representative fields were randomly selected for analysis. The number of cells was quantified using ImageJ software (National Institutes of Health, USA). Briefly, images were duplicated and converted to 8-bit grayscale. A binary mask was created using the Default auto-threshold algorithm with the background set to black. To separate touching or overlapping nuclei, the Watershed command was applied. Finally, particles were analyzed using the Analyze Particles function, counting objects with a size range of 50-500 square pixels. The average number of cells in each area was used for normalizing oxygen consumption rate or extracellular acidification rate value.

**Cleavage Under Targets and Tagmentation (CUT&Tag)**

CUT&Tag experiments were conducted using CUT&Tag Assay Kit following the manufacturer's protocol with slight modification. Briefly, $2.5 \times 10^5$ HU-HSA-2 or HU-HSA-3 cells were prepared for each reaction in duplicate for pan-Kla samples and in singly for H3K4me3, H3K27ac, and RNAPII samples. Cells were washed with 1.0 mL Complete Wash Buffer twice and incubated with activated beads for 5 min at RT, and then antibodies (pan-Kla, H3K4me3, H4K27ac, and phospho-Rpb1) were added and incubated for 1.5 hour at RT. Afterwards, cells were incubated with the secondary antibody (Goat Anti-Rabbit IgG [H+L], 1:50) for 30 min at RT. pAG-Tn5 was introduced to the cells and incubated for 1 hour at RT followed by washing cells with 500 μL Digitonin Buffer twice. DNA antibody binding was tagged by adding magnesium chloride and incubated for 1 hour at 37°C followed by washing cells with 500 μL High Salt Digitonin Buffer twice. To stop tagmentation, 6.75 μL 0.5M EDTA, 8.25 μL 10% SDS and 1.5 μL Proteinase K were added to DNA mixture and proceeded to DNA purification. For DNA amplification, PCR was performed for purified DNA fragments with index primers using PCR Master Mix. The process consists of 72°C for 5 min for 1 cycle, 98°C for 40 sec and 63°C for 10 sec for 13 cycles, and 72°C for 1 min for 1 cycle. Amplified DNA was purified with AMpure. DNA concentration was measured with Qubit using a dsDNA high sensitivity kit. DNA quality was checked with TapeStation using a D5000 High sensitivity kit. DNA libraries were submitted to Rhelixa Co., Ltd. (Tokyo, Japan) and sequenced with the Illumina NovaSeq X Plus (Illumina, CA, USA) to generate a minimum of 20 million paired-end 150 bp reads for pan-Kla samples cultured under 0 mM glucose condition, and 13.3 million paired-end 150 bp reads for pan-Kla cultured in 25 mM glucose condition and H3K4me3, H3K27ac, and RNAPII samples cultured with/without glucose. Sequencing reads that qualified with Fastp v0.26.0 *(52)* were mapped to the CanFam3.1 canine reference genome from Ensembl using Bowtie2 v2.5.4 *(53)*. For sample normalization, sum of fragments coverage was determined by calculating the number of fragments which were ranged from 1 to 1,000 bp in bedpe formats converted from mapped bam files using bedtools v2.31.1 *(54)*. Raw bedgraph files converted from bam files using bamCoverage in the deepTools package v3.5.6 *(55)* were normalized by sum of fragments coverage, then visualized using Integrative Genome Viewer (IGV). For visualizing transcription start sites (TSSs), computeMatrix from deeptools was used with default setting of reference-point mode, then calculated matrix was visualized by plotProfile. For visualizing gene body heatmaps and genomic annotation, plotPeakProf2 and plotAnnopie in ChiPseeker v3.21 were used, respectively *(56, 57)*. To identify significantly enriched peaks in the 0 mM glucose condition relative to the 25 mM condition, peak calling was performed using MACS2 (v2.2.9.1) *(58)*. The 0 mM glucose samples were designated as the treatment (-t), and the 25 mM glucose samples were used as the control (-c). The false discovery rate (FDR) threshold (-q) was set to 0.1 for pan-Kla and H3K4me3 samples, and 0.001 for RNAPII-Ser5 samples. Afterwards, significant peaks were then annotated to genomic features using Homer (v5.1) *(59)* with a custom annotation file generated from the CanFam3.1 canine reference genome (Ensembl). Gene ontology analysis was conducted using PANTHER Pathway in PANTHER v18.0 *(60, 61)*.

**RNA-sequencing**

HU-HSA-2 and HU-HSA-3 were cultured under 0 mM or 25 mM glucose conditions for 48 hours in triplicate. Total RNA was extracted using a NucleoSpin RNA isolation kit according to the manufacturer's instructions. RNA samples were submitted to Rhelixa Co., Ltd. for further analyses. mRNA-seq libraries were constructed using a NEBNext Ultra II Directional RNA Library Prep Kit and sequenced with the Illumina NovaSeq X Plus platform to generate a minimum of 40 million paired-end 150-bp reads. Sequencing reads were mapped to the CanFam3.1 canine reference genome using STAR, and expression levels were estimated using RSEM *(62, 63)*. Differential expression was analyzed with edgeR 3.42.0, and pathway enrichment was evaluated with GSEA v4.4.0 *(64–66)*. To compare transcriptional profiles of our HSA cell lines with other canine cells, we analyzed our mRNA-seq data of HSA cell lines and 29 publicly available canine cell sequence datasets (BioProject accessions PRJNA719562, PRJNA803064, PRJNA590267, PRJNA689618, and PRJNA786902) (Table S2). All FASTQ files were aligned to the CanFam4 reference genome with STAR and gene expression levels were estimated with RSEM. Read counts were analyzed in R v4.3.2. Genes with low CPM were filtered with edgeR (filterByExpr), library sizes were normalized by the trimmed mean of M-values (TMM), and precision weights were estimated with limma-voom. Principal-component analysis was performed on $\log_2$CPM values after removing batch effects using limma 3.56.2. Mean ± SD expression of endothelial markers was summarized per biological group and visualized with ggplot2.

**Reverse transcription quantitative polymerase chain reaction (RT-qPCR)**

Total RNA was extracted with TriPure Isolation Reagent according to the manufacturer's instructions. Reverse transcription was performed using Primescript II 1st strand cDNA Synthesis Kit for 1 µg of total RNA of samples according to the manufacturer's instructions. qPCR samples were prepared using KAPA SYBR FAST qPCR Kit Master Mix (2×) ABI Prism. The reaction solution contained 1× KAPA SYBR FAST qPCR Master Mix, 200 nM forward and reverse primers, 1 µl cDNA and UltraPure DNase/RNase-free distilled water (UPDW). The samples were applied in triplicate and analyzed by StepOne Real-time PCR system. Samples were denatured at 95°C for 20 sec followed by 40 cycles of 95°C for 3 sec and 60°C for 30 sec. RT-qPCR was performed using the primers listed in Table S5. Results were normalized using the geometric mean of reference genes (*RPL32*, *ACTB*, *B2M*, *HMBS*, *TBP* and *YWHAZ*), which were selected from potential internal controls by geNorm *(67)*. UPDW (no template) and no RT samples were used as negative controls and confirmed that no signal was detected for all primer sets. Gene sequences were obtained from Ensembl. Primer3 ver 3.3.0 was used to design primers targeting 80-150 bp products. The primers bind all splice variants and exon-exon junctions. The BLAST database was used to confirm that each primer set did not detect other genes. Primer set specificity was evaluated by checking that each primer set has identical and singular peak in the melting curve. Relative expression levels were calculated by setting the expression levels in the 0 hour samples as 1.

**Single-cell transcriptome analysis**

Tumor tissues were harvested from two PDX models under anesthesia as described in the Animal Study section. Tumor tissues were washed with PBS to remove excess blood and trimmed to carefully remove mouse-derived adipose and connective tissues. The remaining tumor tissue was mechanically dissociated by mincing it into small fragments (~1-2 mm cubes) using sterile scalpels. Tissue fragments were then washed with PBS containing 0.1% bovine serum albumin (BSA) to prevent cell aggregation, and then subjected to two rounds of RBC lysis using an $NH_4Cl$-based buffer with gentle agitation for 5 min. Following a wash with PBS/BSA, the tissue fragments were enzymatically digested in a solution containing 3 mg/mL collagenase I and 1 µg/mL DNase I in DMEM for 50 min at 37°C with intermittent mixing. The digested tissue was gently homogenized by passing through 18G and 23G needles. The resulting cell suspension was then passed through a 40 µm cell strainer to remove any remaining clumps. After another wash and RBC lysis step to ensure purity, dead cells were depleted using the Dead Cell Removal Kit according to the manufacturer's protocol. The number of viable cells was determined with trypan blue staining. The cell concentration was adjusted to approximately 1,000 cells/µL. Five thousand cells per sample were loaded onto a Chromium Controller for single-cell capture. Single-cell gene expression libraries were prepared using the Chromium Next GEM Single Cell 3' Reagent Kits v3.1 for Dual Index following the manufacturer's protocol. The quality and fragment size distribution of the final libraries were assessed using an Agilent Bioanalyzer 2100 with a High Sensitivity DNA Kit. Libraries were then pooled and sequenced on an Illumina NovaSeq 6000 platform with a targeted sequencing depth of 800 million reads per sample.

For data analysis, canine-specific reads were first extracted using XenoCell with default settings *(68)*, and mapping and alignment were conducted by Cell Ranger (10X Genomics). Data were processed with Seurat ver. 4.2.0 using R ver. 4.5.0 in Rstudio ver. 2025.05.0. Metascape analysis for cluster 10 was conductaed using Metascape *(69)*.

**Spatial transcriptome analysis**

Patient information is detailed in Table S1. A tumor tissue sample obtained from a splenectomy was embedded in Optimal Cutting Temperature (OCT) compound, snap-frozen, and stored at -80°C until use. The frozen tissue was sectioned at 10 µm thickness and placed onto a Stereo-seq chip. Stereo-seq library preparation, sequencing, and primary data analysis were performed by AZENTA (Chelmsford, MA, USA). Libraries were sequenced on a DNBSEQ platform, generating 1-1.5 G of paired-end reads, with ensuring >75% of reads achieved a Phred score ≥ Q30. Raw sequencing reads were demultiplexed using the DNBSEQ platform's built-in software, and the quality of the raw data was assessed using fastp v.0.20.0. The final data processing, including alignment to the canine reference genome (ROS_Cfam_1.0) and spatial expression mapping, was performed using the Stereo-seq Analysis Workflow (SAW) pipeline.

**Migration assay**

HU-HSA-2 and HU-HSA-3 were cultured under 0 mM or 25 mM glucose conditions for 24 hours in 6-well plates. Then, the cells were cocultured with RAW264 seeded on ThinCert Cell Culture Inserts for 24 hours. RAW264 cells on the upper surface of the inserts were removed with a cotton swab. Migrated RAW264 on the bottom side of the inserts were fixed with 4% paraformaldehyde for 30 min at RT, and then stained with 0.01% crystal violet for 30 min at RT. The number of cells was counted manually in 10 fields at 200× under a light microscope (BX-41).

**Conditioned medium assay**

HU-HSA-3 cells were cultured in DMEM with or without 25 mM glucose for 48 hours. The supernatant was collected as conditioned medium and used for further analysis after filtering through a 0.2 μm pore filter. D(+)-glucose was added to the conditioned medium obtained from the 0 mM glucose condition to a final concentration of 5.6 mM to allow RAW264 cells to survive. RAW264 cells were cultured with the conditioned medium (glucose 5.6 mM or 25 mM) or complete DMEM (glucose 5.6 mM or 25 mM) for 24 hours. Afterward, total RNA was harvested for RT-qPCR.

**Metabolome analysis**

$1.0 \times 10^5$ HU-HSA-3 cells were seeded in 10 cm dishes with regular DMEM cell culture medium in triplicate and incubated overnight. Next day, cell culture medium was changed to DMEM without glucose and glutamine (#042-32255, FujifilmWako, custom order lacking glutamine) supplemented with 4 mM $^{13}C_5$ L-glutamine. Cells were harvested at 0, 0.5, 3, 24 and 48 hours after changing cell culture medium. Briefly, cells were washed with 3.4% erythritol twice after aspirating cell culture medium. 800 μL methanol and 10 μM internal standard were added, and then extracted solution was centrifuged at 2,300 × g, 4°C for 5 min. Supernatant was collected and ultrafiltered with a 5 kDa cut-off filter at 9,100 × g, 4°C for 3 hours to remove proteins. Samples were then submitted to Human Metabolome Technology (Yamagata, Japan) for further analyses. Metabolic products were analyzed by Agilent CE-TOFMS system (Agilent Technology) *(70)* with fused silica capillary (i.d. 50 μm × 80 cm in total length) in cation and anion modes. The signal to noise ratio in each peak was calculated, and peaks with the ratio more than 3 were used for the analysis. Using mass to charge ratios (m/z) and migration time of each peak, metabolic products were determined based on metabolic product library from Human Metabolome Technology. For quantification of metabolic products, the concentration of total isotope ions in the product was calculated and normalized with the internal standard *(71, 72)*.

**Cell viability assay**

Two thousand cells were seeded in 96-well cell culture plates and cultured in 100 μl DMEM corresponding to each experimental condition. On the next day, cells were treated with either dimethyl sulfoxide (DMSO) or Tunicamycin or Salubrinal each at five different concentrations (10 μg/mL, 1 μg/mL, 0.1 μg/mL, 0.01 μg/mL, and 0.001 μg/mL for Tunicamycin; 100 μM, 10 μM, 1 μM, 0.1 μM, and 0.01 μM for Salubrinal). Survival rates were analyzed using Cell Counting Kit-8 (CCK-8) according to the manufacturer's instructions with slight modifications. Briefly, 10 μl of CCK-8 solution was added to each well 48 hours after adding DMSO or either inhibitor. After 2 hours incubation, 10 μl of 0.1% SDS solution was added to stop the reaction and the absorbance at 450 nm was measured with a microplate reader MTP-320. Survival rates were calculated by setting absorbance of DMSO treated samples as 100%. KyPlot 5.0 software (KyensLab, Inc., Tokyo, Japan) was used to draw survival curves *(73)*.

**Supplementary Figure. 1**

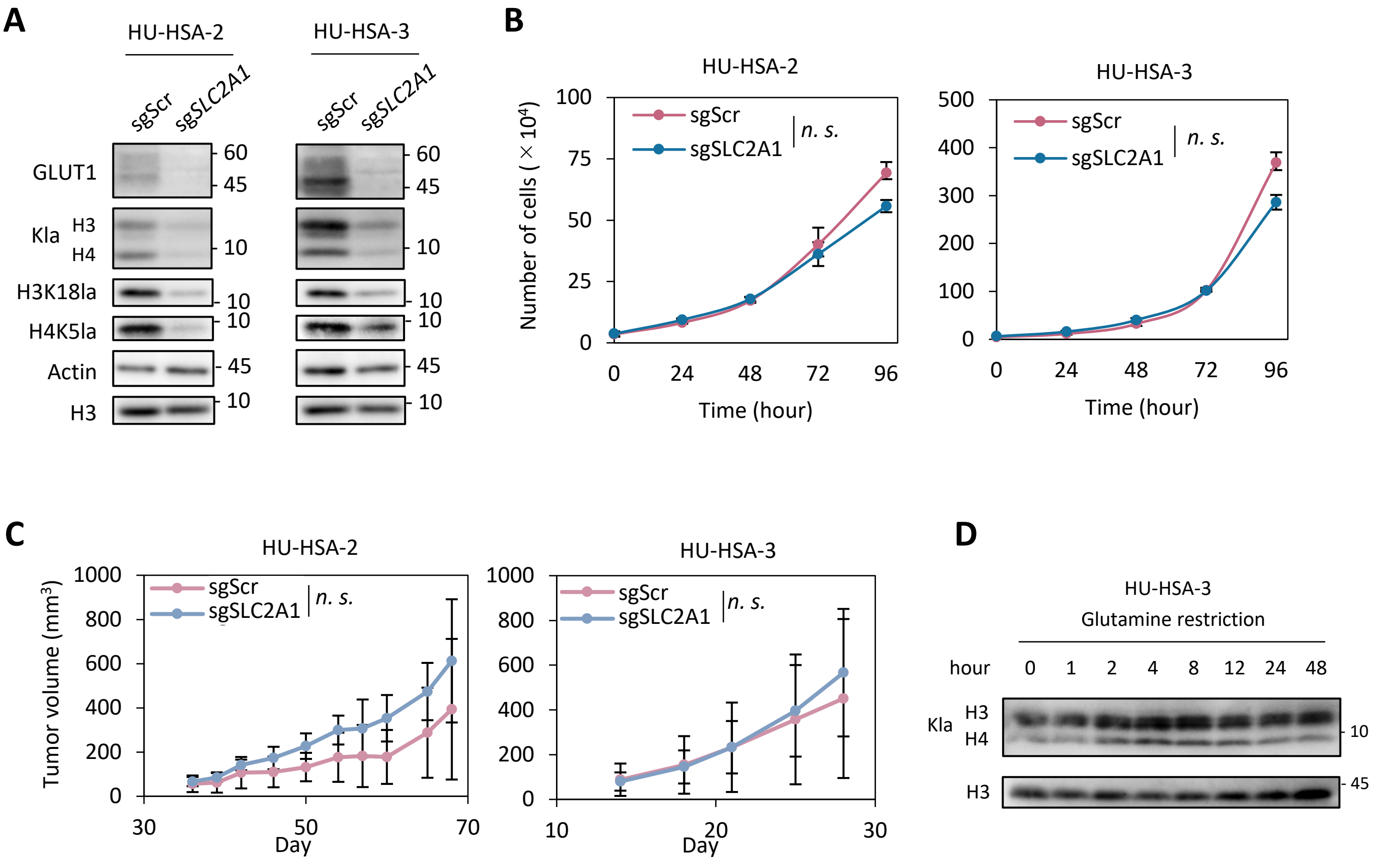


**Fig. S1 *SLC2A1* polyclonal knockout decreases global histone lactylation levels but does not significantly affect HSA cell growth.**

**(A)** Western blot analysis to assess GLUT1 suppression efficiency and global Kla, H3K18la, and H4K5la levels in HSA cell lines expressing the sgScramble (sgScr) and sgSLC2A1. **(B)** *In vitro* growth curves of HSA cell lines expressing sgScr and sgSLC2A1 over 96 hours. Data are presented as average ± SD (three biological replicates). **(C)** *In vivo* tumor growth curves of HSA cell lines expressing sgScr and sgSLC2A1 transplanted subcutaneously into each flank of nude mice Data are presented as average ± SD ($n$ = 6 tumors per group). **(D)** Time-course western blot analysis for Kla in HU-HSA-3 cells cultured in the glutamine-free medium for 0 - 48 hours.
*n.s.*, not significant. Two-way ANOVA.

### Supplementary Figure. 2

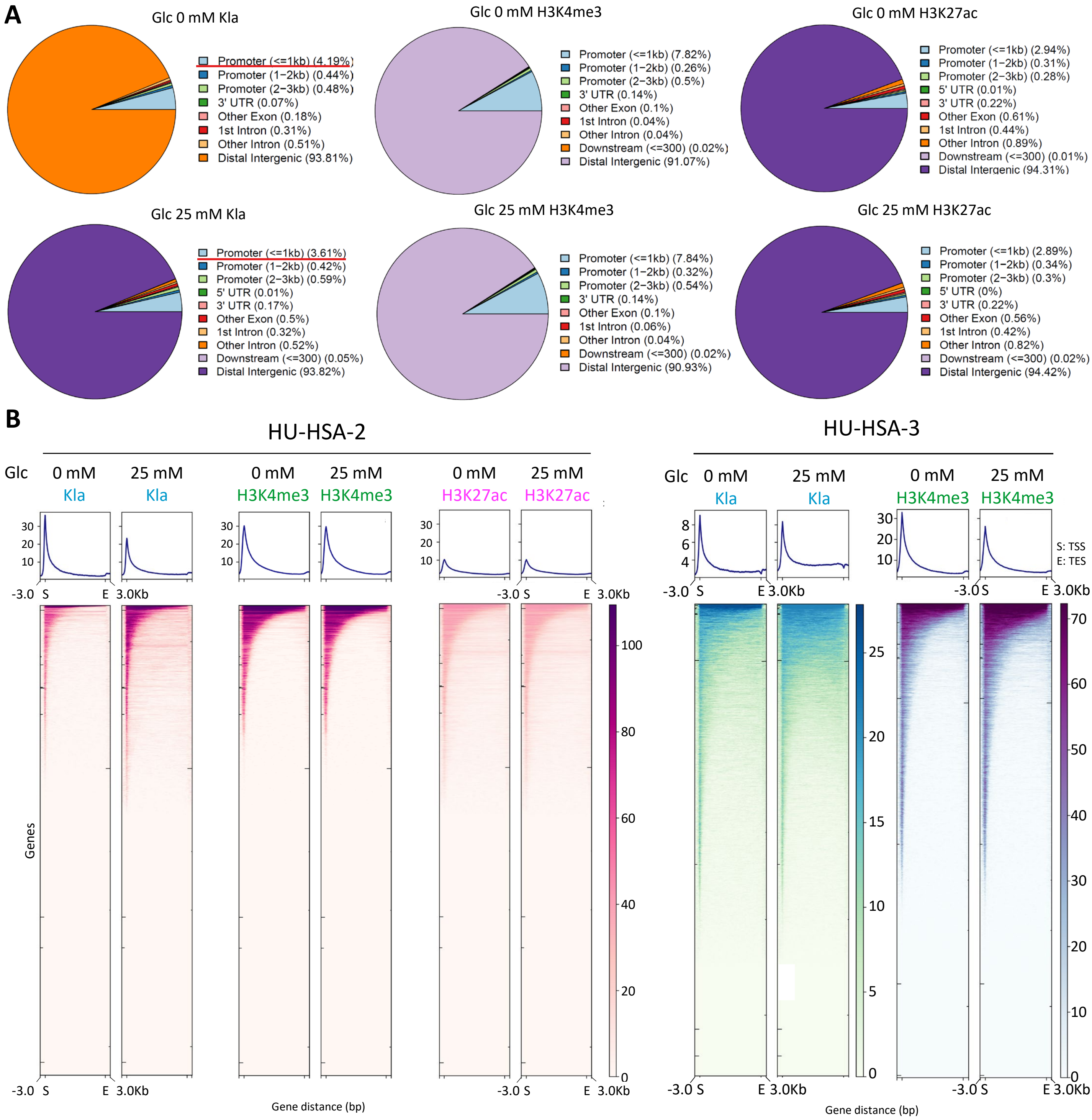


**Fig. S2 Kla is enriched around TSSs under glucose starvation in HSA cells.**

**(A)** Pie charts showing the genomic distribution of Kla, H3K4me3, and H3K27ac enriched peaks obtained from the CUT&Tag analysis in HU-HSA-2 cells cultured with or without glucose for 48 hours. **(B)** Composite profile plots (top) and heatmaps (borrom) on gene-bodies showing the distribution of Kla, H3K4me3, and H3K27ac enrichment in HU-HSA-2 (B) and HU-HSA-3 (C) cells cultured with or without glucose for 48 hours.

### Supplementary Figure. 3

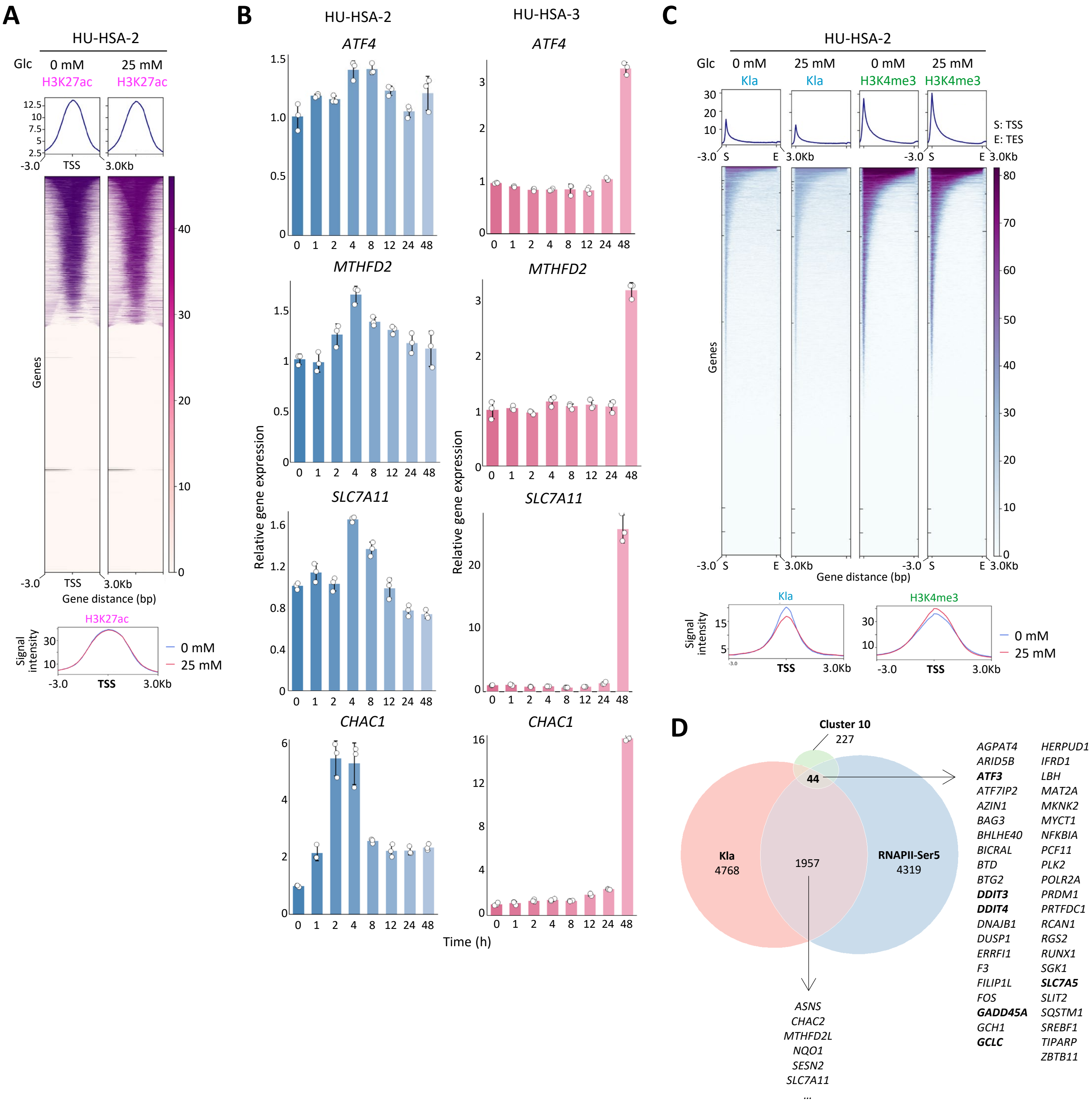


**Fig. S3 Kla is enriched around TSSs of stress response-related genes under glucose starvation in HSA cells.**

**(A)** Composite profile plots (top), heatmaps (middle), and merged profile plots (bottom) around TSS showing the distribution of H3K27ac enrichment in HU-HSA-2 cells cultured with or without glucose for 48 hours. **(B)** Time-course relative expression levels of *ATF4* and selected its target genes in HSA cell lines after glucose starvation. Data are presented as average ± SD from three technical replicates. **(C)** Composite profile plots (top), heatmaps (middle), and profile plots around TSSs (bottom) showing the distribution of Kla and H3K4me3 enrichment in HU-HSA-2 cells cultured with or without glucose for 4 hours. **(D)** Venn diagram showing overlap between cluster 10 marker genes from scRNA-seq and genes with TSSs enrichment of Kla and RNAPII-Ser5 in HU-HSA-3 cells after glucose starvation. 44 of Kla and RNAP-Ser5 enriched genes were cluster 10 markers (listed at right).

### Supplementary Figure. 4

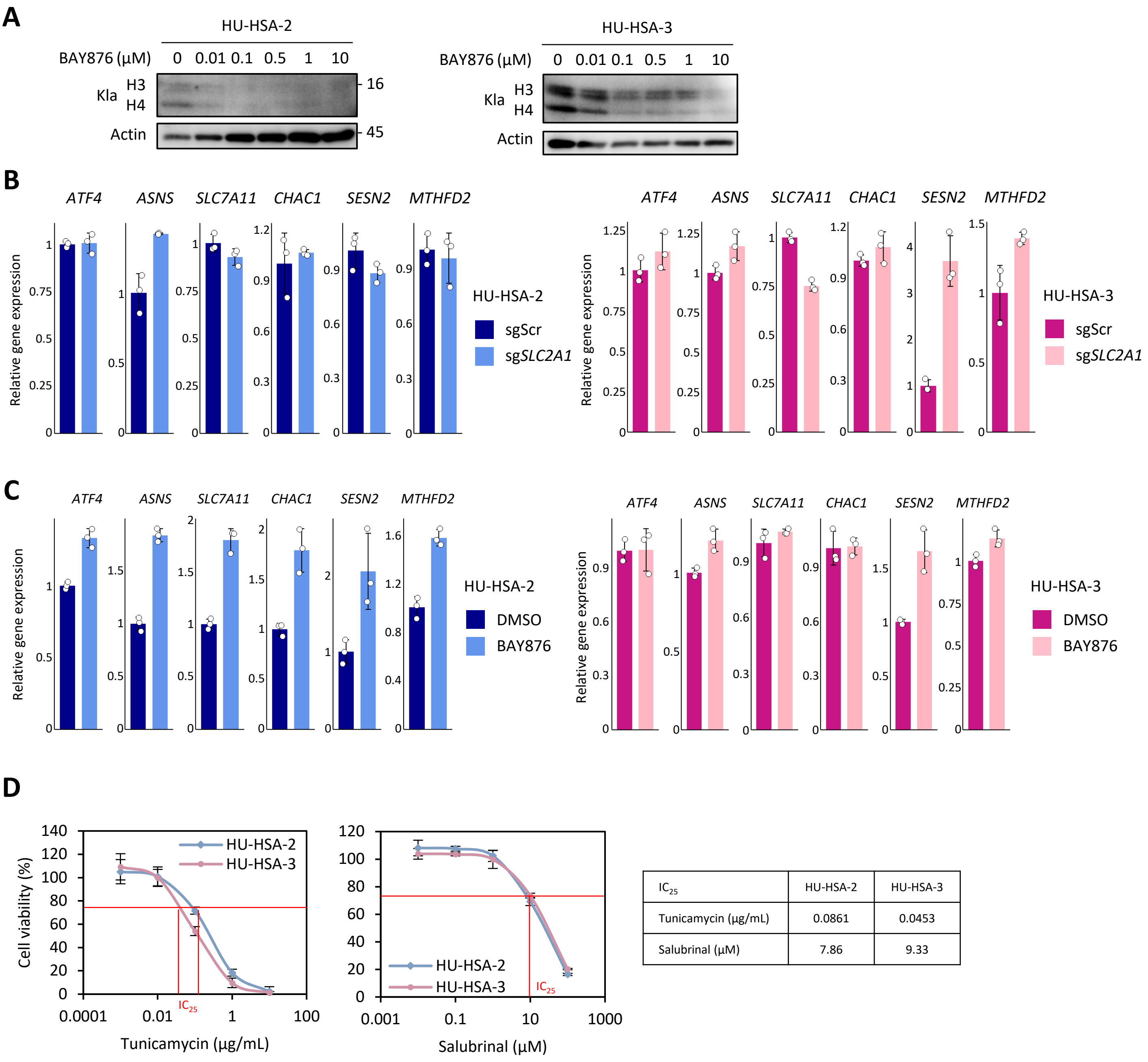


| $IC_{25}$ | HU-HSA-2 | HU-HSA-3 |
|---|---|---|
| Tunicamycin (μg/mL) | 0.0861 | 0.0453 |
| Salubrinal (μM) | 7.86 | 9.33 |

**Fig. S4 GLUT1 inhibition reduces global histone lactylation but fails to activate stress-response genes.**

**(A)** Western blot analysis for Kla in HSA cell lines treated with BAY876 (0 - 10 μM) for 48 hours. **(B, C)** Relative expression levels of key stress response genes in HSA cell lines expressing sgScr or sgSLC2A1 (B), or treated with either DMSO or 0.1 μM BAY876 (C). Data are presented as average ± SD from three technical replicates. Gene expressions were normalized to that of the corresponding controls. **(D)** Cell viability curves of HU-HSA-2 and HU-HSA-3 cells treated with tunicamycin or salubrinal for 48 hours. Red lines indicate the line for 75% survival. Data are presented as average ± SD from three technical replicates.

### Supplementary Figure. 5

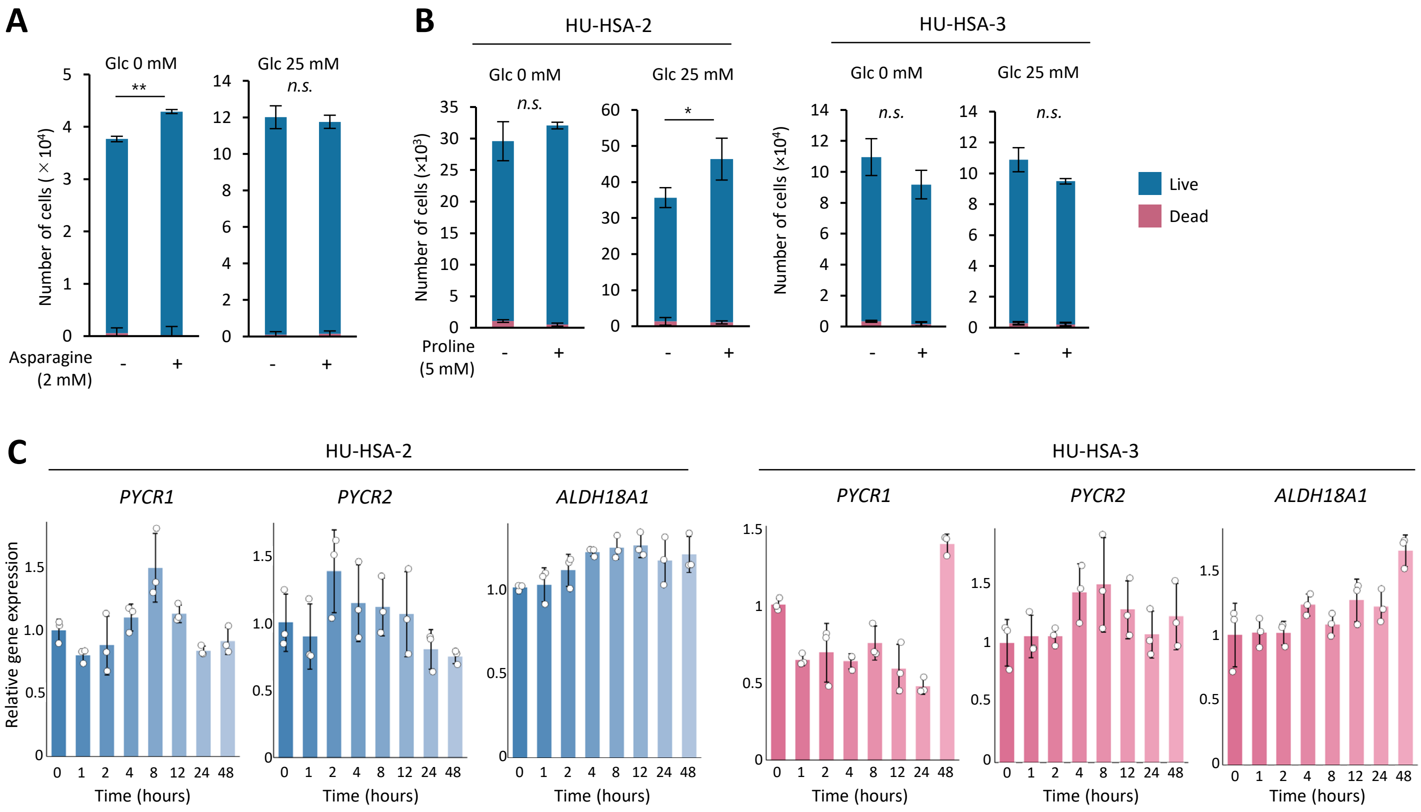


**Fig. S5 Asparagine modestly accelerates HSA cell proliferation, while proline does not.**

**(A)** The number of HU-HSA-2 cells cultured for 72 hours in regular or glucose-free medium supplemented with 2 mM asparagine and 3% FBS. Data are presented as average ± SD from three biological replicates. $^{**}P < 0.01$; n.s., not significant; Student's *t* test. **(B)** The number of HU-HSA-2 and HU-HSA-3 cells cultured for 72 hours in regular or glucose-free medium supplemented with 5 mM proline and 1% FBS. Data are presented as average ± SD from three biological replicates. $^{*}P < 0.05$; n.s., not significant; Student's *t* test. **(C)** Time-course relative expression levels of proline synthesis-related genes in HU-HSA-2 and HU-HSA-3 cells following glucose starvation. Data are presented as average ± SD from three technical replicates.

### Supplementary Figure. 6

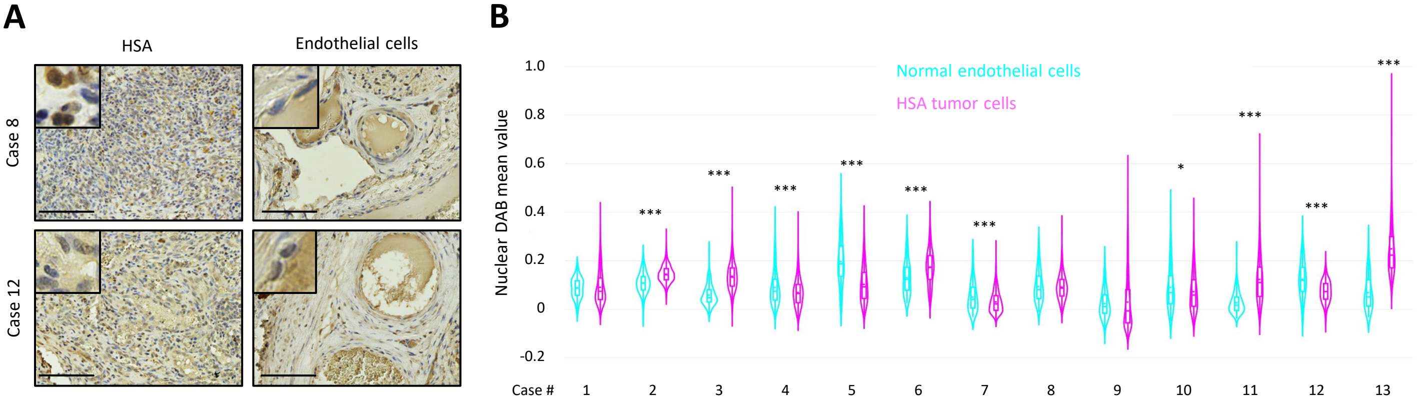


**Fig. S6 IHC analysis for Kla shows no consistent trend between HSA cells and normal endothelial cells.**

**(A)** Representative images of Kla staining in HSA tissues and adjacent normal splenic tissues. Scale bars, 100 µm. **(B)** Violin plots of nuclear pan-Kla intensities in normal endothelial cells versus HSA tumor cells across 13 patient cases. ***$P < 0.001$, *$P < 0.05$. Student's *t* test.

### Table S1 Patient information

**For cell lines and PDX model establishment**

| Patient ID | Breed | Age | Sex | Location |
|---|---|---|---|---|
| HU-HSA-1 | Standard Poodle | 12y3m | M | Spleen |
| HU-HSA-2 | Toy Poodle | 11y1m | M Cast | Spleen |
| HU-HSA-3 | Flat Coated Retriever | 8y | F Spay | Spleen |

**For spatial transcriptomics**

| Breed | Age | Sex | Location |
|---|---|---|---|
| Yorkshire Terrier | 13y3m | M Cast | Spleen |

**For IHC analysis**

| Case number | Breed | Age | Sex | Location |
|---|---|---|---|---|
| 1 | Great Pyrenees | 9y11m | M | Spleen |
| 2 | Miniature Schnauzer | 11y | M | Spleen |
| 3 | Golden retriever | 11y | M | Spleen |
| 4 | Dachshund | 7y9m | M Cast | Spleen |
| 5 | Golden retriever | 7y8m | F Spay | Spleen |
| 6 | Miniature Schnauzer | 12y10m | F Spay | Spleen |
| 7 | Labrador Retriever | 8y | M | Spleen |
| 8 | Mix | 9y3m | F | Spleen |
| 9 | French Bulldog | 13y1m | M | Spleen |
| 10 | Beagle | 12y6m | F Spay | Spleen |
| 11 | Dachshund | 14y5m | M Cast | Spleen |
| 12 | French Bulldog | 7y8m | M Cast | Spleen |
| 13 | Jack Russell Terrier | 9y11m | M Cast | Spleen |

F: female, M: male, Spay: spayed, Cast: castrated

**Table S2 Summary of canine cell sequencing datasets**

| Experiment Accession | Experiment Title | Organism Name | Instrument | Study Accession | Sample Accession |
|---|---|---|---|---|---|
| SRX10509645 | GSM5225598: endothelial cells femoral artery dog A; Canis lupus familiaris; RNA-Seq | Canis lupus familiaris | NextSeq 500 | SRP313354 | SRS8633738 |
| SRX10509646 | GSM5225599: endothelial cells femoral artery dog B; Canis lupus familiaris; RNA-Seq | Canis lupus familiaris | NextSeq 500 | SRP313354 | SRS8633739 |
| SRX10509647 | GSM5225600: endothelial cells femoral artery dog C; Canis lupus familiaris; RNA-Seq | Canis lupus familiaris | NextSeq 500 | SRP313354 | SRS8633740 |
| SRX10509648 | GSM5225601: endothelial cells pulmonary artery dog A; Canis lupus familiaris; RNA-Seq | Canis lupus familiaris | NextSeq 500 | SRP313354 | SRS8633741 |
| SRX10509649 | GSM5225602: endothelial cells pulmonary artery dog B; Canis lupus familiaris; RNA-Seq | Canis lupus familiaris | NextSeq 500 | SRP313354 | SRS8633742 |
| SRX10509650 | GSM5225603: endothelial cells pulmonary artery dog C; Canis lupus familiaris; RNA-Seq | Canis lupus familiaris | NextSeq 500 | SRP313354 | SRS8633743 |
| SRX9779362 | GSM5005184: D17 parental rep 1; Canis lupus familiaris; RNA-Seq | Canis lupus familiaris | Illumina HiSeq 2500 | SRP300293 | SRS7966796 |
| SRX9779363 | GSM5005185: D17 parental rep 2; Canis lupus familiaris; RNA-Seq | Canis lupus familiaris | Illumina HiSeq 2500 | SRP300293 | SRS7966797 |
| SRX9779364 | GSM5005186: D17 parental rep 3; Canis lupus familiaris; RNA-Seq | Canis lupus familiaris | Illumina HiSeq 2500 | SRP300293 | SRS7966798 |
| SRX9779368 | GSM5005190: HMPOS parental rep 1; Canis lupus familiaris; RNA-Seq | Canis lupus familiaris | Illumina HiSeq 2500 | SRP300293 | SRS7966802 |
| SRX9779369 | GSM5005191: HMPOS parental rep 2; Canis lupus familiaris; RNA-Seq | Canis lupus familiaris | Illumina HiSeq 2500 | SRP300293 | SRS7966803 |
| SRX9779370 | GSM5005192: HMPOS parental rep 3; Canis lupus familiaris; RNA-Seq | Canis lupus familiaris | Illumina HiSeq 2500 | SRP300293 | SRS7966804 |
| SRX13340456 | GSM5720812: 1508-1_S1_L001; Canis lupus familiaris; RNA-Seq | Canis lupus familiaris | NextSeq 500 | SRP349649 | SRS11258887 |
| SRX13340457 | GSM5720813: 1508-1_S1_L002; Canis lupus familiaris; RNA-Seq | Canis lupus familiaris | NextSeq 500 | SRP349649 | SRS11258886 |
| SRX13340458 | GSM5720814: 1508-1_S1_L003; Canis lupus familiaris; RNA-Seq | Canis lupus familiaris | NextSeq 500 | SRP349649 | SRS11258888 |
| SRX13340459 | GSM5720815: 1508-1_S1_L004; Canis lupus familiaris; RNA-Seq | Canis lupus familiaris | NextSeq 500 | SRP349649 | SRS11258889 |
| SRX13340460 | GSM5720816: 1508-2_S7_L001; Canis lupus familiaris; RNA-Seq | Canis lupus familiaris | NextSeq 500 | SRP349649 | SRS11258891 |
| SRX13340461 | GSM5720817: 1508-2_S7_L002; Canis lupus familiaris; RNA-Seq | Canis lupus familiaris | NextSeq 500 | SRP349649 | SRS11258890 |
| SRX13340462 | GSM5720818: 1508-2_S7_L003; Canis lupus familiaris; RNA-Seq | Canis lupus familiaris | NextSeq 500 | SRP349649 | SRS11258893 |
| SRX13340463 | GSM5720819: 1508-2_S7_L004; Canis lupus familiaris; RNA-Seq | Canis lupus familiaris | NextSeq 500 | SRP349649 | SRS11258892 |
| SRX13340464 | GSM5720820: 1508-3_S8_L001; Canis lupus familiaris; RNA-Seq | Canis lupus familiaris | NextSeq 500 | SRP349649 | SRS11258895 |
| SRX13340465 | GSM5720821: 1508-3_S8_L002; Canis lupus familiaris; RNA-Seq | Canis lupus familiaris | NextSeq 500 | SRP349649 | SRS11258894 |
| SRX13340466 | GSM5720822: 1508-3_S8_L003; Canis lupus familiaris; RNA-Seq | Canis lupus familiaris | NextSeq 500 | SRP349649 | SRS11258898 |
| SRX13340467 | GSM5720823: 1508-3_S8_L004; Canis lupus familiaris; RNA-Seq | Canis lupus familiaris | NextSeq 500 | SRP349649 | SRS11258896 |
| SRX13340420 | GSM5720800: 1506-1_S3_L001; Canis lupus familiaris; RNA-Seq | Canis lupus familiaris | NextSeq 500 | SRP349649 | SRS11245299 |
| SRX13340421 | GSM5720801: 1506-1_S3_L002; Canis lupus familiaris; RNA-Seq | Canis lupus familiaris | NextSeq 500 | SRP349649 | SRS11245300 |
| SRX13340422 | GSM5720802: 1506-1_S3_L003; Canis lupus familiaris; RNA-Seq | Canis lupus familiaris | NextSeq 500 | SRP349649 | SRS11245301 |
| SRX13340423 | GSM5720803: 1506-1_S3_L004; Canis lupus familiaris; RNA-Seq | Canis lupus familiaris | NextSeq 500 | SRP349649 | SRS11245302 |
| SRX13340448 | GSM5720804: 1506-2_S11_L001; Canis lupus familiaris; RNA-Seq | Canis lupus familiaris | NextSeq 500 | SRP349649 | SRS11245307 |
| SRX13340449 | GSM5720805: 1506-2_S11_L002; Canis lupus familiaris; RNA-Seq | Canis lupus familiaris | NextSeq 500 | SRP349649 | SRS11245308 |
| SRX13340450 | GSM5720806: 1506-2_S11_L003; Canis lupus familiaris; RNA-Seq | Canis lupus familiaris | NextSeq 500 | SRP349649 | SRS11245309 |
| SRX13340451 | GSM5720807: 1506-2_S11_L004; Canis lupus familiaris; RNA-Seq | Canis lupus familiaris | NextSeq 500 | SRP349649 | SRS11245310 |
| SRX13340452 | GSM5720808: 1506-3_S6_L001; Canis lupus familiaris; RNA-Seq | Canis lupus familiaris | NextSeq 500 | SRP349649 | SRS11245311 |
| SRX13340453 | GSM5720809: 1506-3_S6_L002; Canis lupus familiaris; RNA-Seq | Canis lupus familiaris | NextSeq 500 | SRP349649 | SRS11245312 |
| SRX13340454 | GSM5720810: 1506-3_S6_L003; Canis lupus familiaris; RNA-Seq | Canis lupus familiaris | NextSeq 500 | SRP349649 | SRS11245313 |
| SRX13340455 | GSM5720811: 1506-3_S6_L004; Canis lupus familiaris; RNA-Seq | Canis lupus familiaris | NextSeq 500 | SRP349649 | SRS11245314 |
| SRX7177537 | GSM4175965: UUA - uninfected cells from uninfected well; Canis lupus familiaris; RNA-Seq | Canis lupus familiaris | Illumina NovaSeq 6000 | SRP230456 | SRS5684513 |
| SRX7177538 | GSM4175966: UUB - uninfected cells from uninfected well; Canis lupus familiaris; RNA-Seq | Canis lupus familiaris | Illumina NovaSeq 6000 | SRP230456 | SRS5684514 |
| SRX7177539 | GSM4175967: UUC - uninfected cells from uninfected well; Canis lupus familiaris; RNA-Seq | Canis lupus familiaris | Illumina NovaSeq 6000 | SRP230456 | SRS5684515 |
| SRX7177540 | GSM4175968: UUD - uninfected cells from uninfected well; Canis lupus familiaris; RNA-Seq | Canis lupus familiaris | Illumina NovaSeq 6000 | SRP230456 | SRS5684516 |
| SRX14030411 | RNA-Seq of Canis lupus familiaris: Fibroblast | Canis lupus familiaris | Illumina MiSeq | SRP358051 | SRS11866026 |
| SRX14030410 | RNA-Seq of Canis lupus familiaris: Fibroblast | Canis lupus familiaris | Illumina MiSeq | SRP358051 | SRS11866024 |
| SRX14030407 | RNA-Seq of Canis lupus familiaris: Fibroblast | Canis lupus familiaris | Illumina MiSeq | SRP358051 | SRS11866022 |
| SRX14030406 | RNA-Seq of Canis lupus familiaris: Fibroblast | Canis lupus familiaris | Illumina MiSeq | SRP358051 | SRS11866021 |

**Table S3 Antibodies for IHC, WB and CUT&Tag**

| Antibody | Supplier | Catalog number | Clone name | Dilution |
|---|---|---|---|---|
| anti-CD31 | Abcam | ab134168 | EP3095 | IHC 1:200<br>WB 1:1000 |
| anti-Von Willebrand Factor | Agilent | A0082 | | IHC 1:500<br>WB 1:1000 |
| anti-VEGF Receptor 2 (KDR) | Abcam | ab2349 | | WB 1:1000 |
| anti-L-Lactyl-Histone H3 Lys18 rabbit monoclonal antibody | PTM Biolabs | PTM-1406RM | | IHC 1:250<br>WB 1:1000 |
| anti-Iba1 | Fujifilm Wako | 019-19741 | | IHC 1:500 |
| anti-CD3 | Agilent Technologies | IR503 | | IHC ready to use |
| anti-CD204 | Medicinal Chemistry Pharmaceutical Co., Ltd. | KT022 | SRA-E5 | IHC 1:200 |
| anti-L-Lactyllysine | PTM Biolabs | PTM-1401RM | 9H1L6 | C&T 1:50<br>IHC 1:200 |
| anti-Tri-Methyl-Histone H3 (Lys4) | Cell Signaling Technology | 9751 | C42D8 | C&T 1:50 |
| anti-Acetyl-Histone H3 (Lys27) | Cell Signaling Technology | 8173 | D5E4 | C&T 1:100 |
| anti-Phospho-Rpb1 CTD (Ser5) | Cell Signaling Technology | 13523 | D9N5I | C&T 1:50 |
| anti-L-Lactyl-Histone H4 (Lys5) Rabbit mAb | PTM Biolabs | PTM-1407RM | | WB 1:1000 |
| anti-Acetylated Histone H3 | Active Motif | 39040 | | WB 1:5000 |
| anti-Acetylated Histone H4 | Santa Cruz Biotechnology, Inc. | sc-377520 | E-5 | WB 1:500 |
| anti-Actin | Sigma-Aldrich | MAB1501 | C4 | WB 1:10000 |
| anti-Histone H3 | MAB Institute | MABI0001-20 | CMA301 | WB 1:25000 |
| anti-ATF-4 | Santa Cruz Biotechnology, Inc. | sc-390063 | B-3 | WB 1:1000 |
| anyi-Asparagine synthethase | Santa Cruz Biotechnology, Inc. | sc-365809 | G-10 | WB 1:1000 |
| anti-Glut1 | Santa Cruz Biotechnology, Inc. | sc-377228 | A-4 | WB 1:1000 |
| Goat anti-Mouse IgG (H+L) | Thermo Fisher Scientific | G21040 | | WB 1:10000 |
| Goat anti-Rabbit IgG (H+L) | Thermo Fisher Scientific | G21234 | | WB 1:10000 |

**Table S4 Regents and instruments**

| | Product name | Supplier | Catalog number |
|---|---|---|---|
| Reagent | Canine genotypes panel 2.1 | Thermo Fisher Scientific, MA, USA | F864S |
| | CnAOEC | Cell Applications, CA,USA | Cn304-05 |
| | Dulbecco's Modified Eagle Medium with High Glucose | Fujifilm Wako Pure Chemical Industries, Osaka, Japan | 044-29765 |
| | Dulbecco's Modified Eagle Medium with no Glucose | Fujifilm Wako Pure Chemical Industries | 042-32255 |
| | Dulbecco's Modified Eagle Medium with no glutamine | Fujifilm Wako Pure Chemical Industries | 045-30285 |
| | Fetal bovine serum | Gibco, NY, USA | 10270-106 |
| | penicillin–streptomycin solution | Fujifilm Wako Pure Chemical Industries | 168-23191 |
| | EASYstrainer (70 μm) | Greiner Bio-One, Kremsmünster, Austria | 542070 |
| | TaKaRa BCA Protein Assay Kit | Takara Bio, Kusatsu, Japan | T9300A |
| | Immobilon-P transfer membranes | Merck Millipore, MA, USA | IPVH00010 |
| | Can Get Signal Solution | TOYOBO, Osaka, Japan | NKB-101 |
| | Immobilon Western Chemiluminescent HRP substrate | Merck Millipore | WBKLS0500 |
| | 10% normal goat serum | Nichirei biosciences, Tokyo, Japan | 426042 |
| | goat anti-rabbit IgG conjugated peroxidase | Nichirei biosciences | 414341 |
| | 3.3'-diaminobenzidine | Dojindo, Kumamoto, Japan | 349-00903 |
| | goat anti-rabbit IgG conjugated alkaline phosphatase | Nichirei biosciences | 414251 |
| | goat anti-mouse IgG conjugated alkaline phosphatase | Nichirei biosciences | 414241 |
| | New Fuchsin solution | Nichirei biosciences | 415161F |
| | 12-well plate | Greiner Bio-One | 665180 |
| | Asparagine | MP Biomedicals, CA, USA | 590-20432 |
| | Proline | Fujifilm Wako Pure Chemical Industries | 163-04601 |
| | 0.25w/v% Trypsin-1mmol/l EDTA･4Na Solution with Phenol Red | Fujifilm Wako Pure Chemical Industries | 201-16945 |
| | Lipofectamine 3000 | Thermo Fisher Scientific | L3000015 |
| | 0.45 μm pore filter | Sartorius, Göttingen, Germany | S7598FXOSK |
| | Domitor (medetomidine) | ZENOAQ, Tokyo, Japan | N/A |
| | Dormicum (midazolam) | Maruishi Pharmaceutical Co., Ltd. Osaka, Japan | 211-762100 |
| | Vetorphale (butorphanol) | Meiji Seika Pharma Co., Ltd. Tokyo, Japan | N/A |
| | Atipame (atipamezole) | Kyoritsu Seiyaku Corporation, Tokyo, Japan | N/A |
| | Seahorse XFp Cell Culture Miniplate | Agilent Technology, CA, USA | 103025-100 |
| | Seahorse XF base medium without phenol red | Agilent Technology | 103335-100 |
| | L-glutamine | Fujifilm Wako Pure Chemical Industries | 073-05391 |
| | D(+)-glucose | Fujifilm Wako Pure Chemical Industries, | 047-31161 |
| | ATP rate assay Kit | Agilent Technology | 103591-100 |
| | mitochondrial oxidation assay Kit | Agilent Technology | 103270-100 |
| | Hoechst 33342 | Dojindo, Kumamoto, Japan | 346-07951 |
| | CUT&Tag Assay Kit | Cell Signaling Technology, MA, USA | 77752 |
| | PCR Master Mix | New England Biolabs, MA, USA | M0541S |
| | AMpure | Beckman Coulter, CA, USA | BC-A63880 |
| | dsDNA high sensitivity kit | Invitrogen, MA, USA | Q33230 |
| | D5000 High sensitivity kit | Agilent Technology | 5067-5089 |
| | NucleoSpin RNA isolation kit | Macherey-Nagel GmbH & Co. Düren, Germany | 740955.5 |
| | TriPure Isolation Reagent | Roche, Basel, Switzerland | 11667157001 |
| | Primescript II 1st strand cDNA Synthesis Kit | Takara Bio | 6210 |
| | KAPA SYBR FAST qPCR Kit Master Mix (2×) ABI Prism | KAPA Biosystems, MA, USA | KK4605 |
| | ThinCert Cell Culture Inserts | Greiner Bio-One | 657630 |
| | 0.2 μm pore filter | Sartorius | S7597FXOSK |
| | Dead Cell Removal Kit | Miltenyi Biotec, Bergisch Gladbach, Germany | 130-090-101 |
| | Chromium Next GEM Single Cell 3′ Kit v3.1, 4 rxns | 10x Genomics, CA, USA | PN-1000269 |
| | Chromium Next GEM Chip G Single Cell Kit, 16 rxns | 10x Genomics | PN-1000127 |
| | Dual Index Kit TT Set A, 96 rxns | 10x Genomics | PN-1000215 |
| | High Sensitivity DNA Kit | Agilent Technology | 5067-4727 |
| | Tissue-Tek® O.C.T. Compound | Sakura Finetek Japan Co.,Ltd., Tokyo, Japan | 4583 |
| | 13C5 L-glutamine | Fujifilm Wako Pure Chemical Industries, Osaka, Japan | CLM-1822-H-0.1 |
| | erythritol | Tokyo Chemical Industry Co., Ltd., Tokyo, Japan | E0021 |
| | Ultracentiifugation for 5 kDa cut-off filter | Human Metabolome Technology, Tsuruoka, Japan | UFC3LCCNB HMT |
| | 96 well cell culture plates | Greiner Bio-One | 655180 |
| | Cell Counting Kit-8 | Dojindo | 343-07623 |
| | Eukitt | ORSAtec, Bobingen, Germany | 6.00.01.0001.06.01.EN |
| Instruments | BRANSON Sonifier 450 | Branson Ultrasonics Corporation, CT, USA | |
| | Image Quant LAS 4000 mini luminescent image analyzer | Cytiva, MA, USA | |
| | NanoZoomer 2.0-RS | Hamamatsu Photonics, Hamamatsu, Japan | |
| | CellDrop BF | DeNovix, DE, USA | |
| | Seahorse XFp Analyzer | Agilent Technology | |
| | EVOS FL | Thermo Fisher Scientific | |
| | Qubit | Invitrogen, MA, USA | |
| | TapeStation | Agilent Technology | |
| | StepOne Real-time PCR system | Thermo Fisher Scientific | |
| | BX-41 | Olympus, Tokyo, Japan | |
| | Chromium Controller | 10x Genomics | |
| | MTP-320 | Corona Electric Co.,Ltd, Ibaraki, Japan | |
| | Tissue-Tek VIP5 Jr | Sakura Finetek Japan Co.,Ltd | |

**Table S5 qPCR primer sequences**

| Species | Target | Sequence (Forward) | Sequence (Reverse) | Gene ID |
|---|---|---|---|---|
| Canine | *RPL32* | TGGTTACAGGAGCAACAAGAAA | GCACATCAGCAGCACTTCA | ENSCAFG00000004821 |
| | *TBP* | ATAAGAGAGCCCCGAACCAC | TTCACATCACAGCTCCCCAC | ENSCAFG00000004119 |
| | *YWHAZ* | CGAAGTTGCTGCTGGTGA | TTGCATTTCCTTTTTGCTGA | ENSCAFG00000000580 |
| | *ACTB* | CCAGCAAGGATGAAGATCAAG | TCTGCTGGAAGGTGGACAG | ENSCAFG00000016020 |
| | *HMBS* | TCACCATCGGAGCCATCT | GTTCCCACCACGCTCTTCT | ENSCAFG00000012342 |
| | *B2M* | ACGGAAAGGAGATGAAAGCA | CCTGCTCATTGGGAGTGAA | ENSCAFG00000013633 |
| | *PECAM1* | AACTTCACCATCCAGAAGG | TCCACTGGGGCTATCACC | ENSCAFG00000011740 |
| | *VWF* | AAGCAGACGATGGTGGATTC | AATGTCCAGGAATGGCTCAG | ENSCAFG00000015228 |
| | *KDR* | GGTATGGTCCTTGCCTCAGA | CAGTGGTATCCGTGTCATCG | ENSCAFG00000002079 |
| | *ATF4* | CTTAAGCCATGGCGCTTTTC | GGAATGTGCTTAATTCGAAGGTG | ENSCAFG00000001324 |
| | *ASNS* | TTGGGTTTTGTGCCACCATG | AGAAAGGAAGAGGGGAAAGCTG | ENSCAFG00000002222 |
| | *SLC7A11* | ATTCATGTCCGCAAGCACAC | TGCCAGCCCAATAAAAAGCC | ENSCAFG00000003749 |
| | *SESN2* | TTAGCTGCTTTTGGCGTCTG | TGCAGAAACTCAGCCATGTG | ENSCAFG00000011842 |
| | *CHAC1* | AGATCATGAGGGCTGCACTTG | TAGCCGCCAAGTACTGCTTC | ENSCAFG00000009414 |
| | *DDIT3* | GCGGATCATGTTGAAGATGAGC | TCAGCTGCCATCTCTACAGTTG | ENSCAFG00000030112 |
| | *MTHFD2* | TGTAGATGGCCTCCTTGTTCAG | AACAGCGTTGCAGACCTTTC | ENSCAFG00000008716 |
| | *PYCR1* | GCCACACATCATCCCCTTTATC | AACGCCATCAGCTTCTTCTC | ENSCAFG00000005906 |
| | *PYCR2* | TGTCGGCTCACAAGATCATAGC | TCACCGTCTCCTTGTTGTTCC | ENSCAFG00000016140 |
| | *ALDH18A1* | ATGGAAGCCAAGGTGAAAGC | TGGGTTCCGTTGGCAATAAC | ENSCAFG00000008339 |

**Table S6 sgRNA sequences**

| Species | Target | sgRNA sequence |
|---|---|---|
| Canine | sgScr1 | ATTCTCTCGACATCTGGTGG |
| | sgScr2 | GCTCGGTAACTAACCGGTGC |
| | sg*SLC2A1* | AGTGTTGTAGCCAAACTGCA |
| | sg*ATF4* -1 | TCCAGTAAAGTCCCGCGACA |
| | sg*ATF4* -2 | TTGGTCAGTGCCTCAGACAA |
